# *Mycobacterium tuberculosis* complex lineage 5 exhibits high levels of within-lineage genomic diversity and differing gene content compared to the type strain H37Rv

**DOI:** 10.1101/2020.06.22.164186

**Authors:** C. N’Dira Sanoussi, Mireia Coscolla, Boatema Ofori-Anyinam, Isaac Darko Otchere, Martin Antonio, Stefan Niemann, Julian Parkhill, Simon Harris, Dorothy Yeboah-Manu, Sebastien Gagneux, Leen Rigouts, Dissou Affolabi, Bouke C. de Jong, Conor J. Meehan

## Abstract

Pathogens of the *Mycobacterium tuberculosis* complex (MTBC) are considered monomorphic, with little gene content variation between strains. Nevertheless, several genotypic and phenotypic factors separate the different MTBC lineages (L), especially L5 and L6 (traditionally termed *Mycobacterium africanum*), from each other. However, genome variability and gene content especially of L5 and L6 strains have not been fully explored and may be potentially important for pathobiology and current approaches for genomic analysis of MTBC isolates, including transmission studies.

We compared the genomes of 358 L5 clinical isolates (including 3 completed genomes and 355 Illumina WGS (whole genome sequenced) isolates) to the L5 complete genomes and H37Rv, and identified multiple genes differentially present or absent between H37Rv and L5 strains. Additionally, considerable gene content variability was found across L5 strains, including a split in the L5.3 sublineage into L5.3.1 and L5.3.2. These gene content differences had a small knock on effect on transmission cluster estimation, with clustering rates influenced by the selection of reference genome, and with potential over-estimation of recent transmission when using H37Rv as the reference genome.

Our data show that the use of H37Rv as reference genome results in missing SNPs in genes unique for L5 strains. This potentially leads to an underestimation of the diversity present in the genome of L5 strains and in turn affects the transmission clustering rates. As such, a full capture of the gene diversity, especially for high resolution outbreak analysis, requires a variation of the single H37Rv-centric reference genome mapping approach currently used in most WGS data analysis pipelines. Moreover, the high within-lineage gene content variability suggests that the pan-genome of *M. tuberculosis* is at least several kilobases larger than previously thought, implying a concatenated or reference-free genome assembly (*de novo*) approach may be needed for particular questions.

**Data summary:** Sequence data for the Illumina dataset are available at European Genome-phenome Archive (EGA; https://www.ebi.ac.uk/ega/) under the study accession numbers PRJEB38317 and PRJEB38656. Individual runs accession numbers are indicated in Table S8.

PacBio raw reads for the L5 Benin genome are available on the ENA accession SAME3170744. The assembled L5 Benin genome is available on NCBI with accession PRJNA641267. To ensure naming conventions of the genes in the three L5 genomes can be followed, we have uploaded these annotated GFF files to figshare at https://doi.org/10.6084/m9.figshare.12911849.v1.

Custom python scripts used in this analysis can be found at https://github.com/conmeehan/pathophy.

## Introduction

Tuberculosis (TB) is caused by pathogenic bacteria of the *Mycobacterium tuberculosis* complex (MTBC) that consists of 9 human-adapted lineages and several animal-adapted lineages (Mireia Coscolla and Gagneux 2014; Ngabonziza et al. 2020; M Coscolla et al. 2020; Brites et al. 2018). This group is highly clonal with no horizontal gene transfer (Boritsch et al. 2016; Chiner-Oms et al. 2019). Strains of particular lineages are primarily defined by large sequence polymorphisms (LSP, the presence or deletion of genomic regions) such as the TbD1 region (MTBC specific deletion 1), other regions of difference (RD) (Boritsch et al. 2016; Brosch et al. 2002; Gordon et al. 1999; Behr et al. 1999; Gagneux et al. 2006), and signature single nucleotide polymorphisms (SNPs)(Coll et al. 2014). RDs are particular long sequence genomic region deleted in some groups of MTBC strains but present in others; so that MTBC strains can be classified in groups using that LSP (Gordon et al. 1999; Brosch et al. 2002; Mostowy et al. 2002; Gagneux et al. 2006). Broadly, the lineages of the MTBC occur within three major clades: (1) L1-L4 and L7 form one group, (2) L5, L6, L9 and the animal lineages form another, and (3) L8 sits within its own clade (Coscolla and Gagneux 2014; Ngabonziza et al. 2020; Coscolla et al. 2020), based on the presence/absence of specific RDs, especially TbD1 (Firdessa et al. 2013; Brosch et al. 2002; Nebenzahl-Guimaraes et al. 2016). L5 and L6 are also called *M. tuberculosis* var. africanum (or historically *M. africanum* West-african 1 and 2 respectively) (Riojas et al. 2018; Niemann et al. 2004) and are restricted to West Africa where they cause up to 40% of human TB (de Jong, Antonio, and Gagneux 2010). Reasons for the geographical restriction of L5 and L6 remain unclear, however, adaptation to particular human subpopulation has been suggested (Intemann et al. 2009; Thye et al. 2011; Comas et al. 2013; Gagneux et al. 2006).

Several phenotypic and genotypic features separate L5 and L6 from the other human-adapted lineages. Some TB diagnostics have a lower performance for L5 and L6 strains, compared to other MTBC lineages (Sanoussi et al. 2018; Ofori-Anyinam et al. 2016) and these lineages are less likely to grow in culture (Sanoussi et al. 2017), with dysgonic appearance on solid medium (Leao et al. 2004; Pattyn et al. 1970; Sanoussi et al. 2018; Niemann et al. 2004). Mutations in genes essential for growth in culture were identified for L6 strains (Gehre et al. 2013), yet for L5 strains the reasons for the difficulty in growth remain unclear but may be related to the presence of the TbD1 region and the absence of RD9 (Brosch et al. 2002). L6 strains and those of the closely related animal strains/lineages such as *M. bovis* are reported to be less virulent in humans than those from other human-adapted MTBC lineages in population-based studies (de Jong et al. 2008; Magnus 1966; Reiling et al. 2013) and in genome studies of mutations in virulence regulation genes/systems (Gonzalo-Asensio et al. 2014; Ofori-Anyinam et al. 2020). Infection with an L6 strain progresses slowly to TB disease, and is associated with impaired immunity in some settings but not all (e.g. HIV infection (de Jong et al. 2008)). In contrast, little is known about the genomics, virulence and disease progression of L5 strains.

Whole genome sequencing (WGS) based on next generation sequencing (NGS) of MTBC strains often involves data analysis by mapping of short sequence reads to a reference genome, usually H37Rv (Cole et al. 1998; Camus et al. 2019; Meehan et al. 2019) or a reconstructed ancestor with the same gene content as H37Rv (Comas et al. 2013; Goig et al. 2019, 2020). The resulting SNPs are then used for drug resistance determination, subtyping and transmission analyses (Meehan et al. 2019). However, since H37Rv is a L4 strain, it might not be representative of other MTBC lineages, including L5 strains. Thus, if H37Rv is used as basis for genome analysis, several genes or larger genomic regions may be missed resulting in an underestimation of genome diversity among strains of other lineages that may, however, provide additional information e.g. for transmission analysis.

The members of the MTBC evolved from an environmental organism to an obligate pathogen through genome reduction and acquisition of new genes (Gagneux 2018) and it is known that some differences in gene content exist between lineages (Meehan et al. 2019; Bifani et al. 2000; Periwal et al. 2015; Kato-Maeda et al. 2001; O’Toole and Gautam 2017; Gagneux et al. 2006; Tsolaki et al. 2005). Furthermore, it was reported that genes had a higher genetic diversity among L6 strains compared to L5 ones (Darko Otchere et al. 2018), but the difference in gene content between these two lineages was not investigated. Also, little is known about the gene differences between L4 and L5 and the potential limitations this may impose for in-depth analysis of L5 genome studies (e.g. sub-lineage detection and transmission tracking). Similarly, little is known about within-lineage diversity in terms of gene content.

To address these questions, we assessed the gene content diversity of L5 by analyzing in the context of whole genome sequencing and reference selection. To this end, we analyzed 358 genomes of L5 strains including three completed genomes and compared them to H37Rv and closely related lineages (L6, *M. tuberculosis* var. bovis (hereafter called *M. bovis*)(Riojas et al. 2018)). Our main focus was to determine particularities in the genomes of L5 strains and to define the level of within-lineage gene content differences among strains of this lineage of the MTBC.

## Materials and Methods

### Genomes

#### Complete genomes (PacBio, completed long reads)

Three completed L5 genomes from clinical isolates, all sequenced with the PacBio SMRT technology, were analyzed. One genome was from a Benin isolate (PcbL5Ben; sequenced in this study), one from an isolate from The Gambia (PcbL5Gam, WBB1453_11-00429-1) (Phelan et al. 2018), and one from Nigeria (PcbL5Nig, WBB1454_IB091-1) (Phelan et al. 2018). These genomes are also from disparate parts of the L5 diversity, representing a broad range of L5 sub-lineages, as were recently defined (Coscolla et al. 2020). The previously published reference/complete genomes H37Rv (L4) (NC_000962.3)(Cole et al. 1998; Camus et al. 2019; Lew et al. 2011), L6 (GM041182, Genbank accession n°: FR878060.1, GCF_0001593225.1_ASM159322v1) (Bentley et al. 2012) and *M. bovis* (AF2122/97, accession: LT708304.1) (Malone et al. 2017) were also included.

#### Whole genomic DNA extraction

Genomic DNA extraction was performed on growth from fresh Löwenstein-Jensen slants using the semi-automated Maxwell 16 Cell DNA purification kit in the Maxwell 16 machine, or from the late exponential phase of growth in 7H9 medium, using the CTAB method (Belisle and Sonnenberg 1998; Coscolla et al. 2020)

#### Assembly and annotation of the completed PacBio genomes

The Benin Pacbio genome was assembled using HGAP (Chin et al. 2013) and Quiver (Chin et al. 2013) and checked for sufficient quality and coverage (a 10,000 bp sliding window coverage was always above 65x and average coverage across entire genome is 107x). The genome sequences of all three completed PacBio genomes were annotated using Prokka-1.12 (Seemann 2014) based on the reference genome H37Rv annotation.

### Gene presence-absence analysis

Gene content differences were assessed with an all-vs-all BLASTn approach, using BLAST+ version+2.8.1 (Altschul et al. 1990; Camacho et al. 2009) (Fig.1). For a specific genome-genome comparison, the following procedure was used: the gene sequences of genome 1 (ffn file; coding regions only) were compared to those of genome 2 using the default cut-offs to look for any homology for each gene. Those genes found in genome 1 and not in genome 2 were then compared to the completed PacBio genome (fna file; coding and non-coding regions) of genome 2 to look for pseudogenes (herein qualified as “suspected pseudogenes”) using BLASTn. These “suspected pseudogenes” were then confirmed using tBLASTn of the genome 1 protein sequences (faa file) compared to the completed PacBio genome (fna file) of genome 2. This procedure was used to compare all three L5 completed PacBio genomes to each other as well as each to H37Rv. Those genes present or absent in H37Rv (or pseudogenes in either) were compared in a similar manner to the L6 and *M. bovis* reference strains to determine if these genes/pseudogenes are L5-specific (i.e. present in L5, but not in H37Rv, L6 and *M. bovis*).

**Figure 1.**
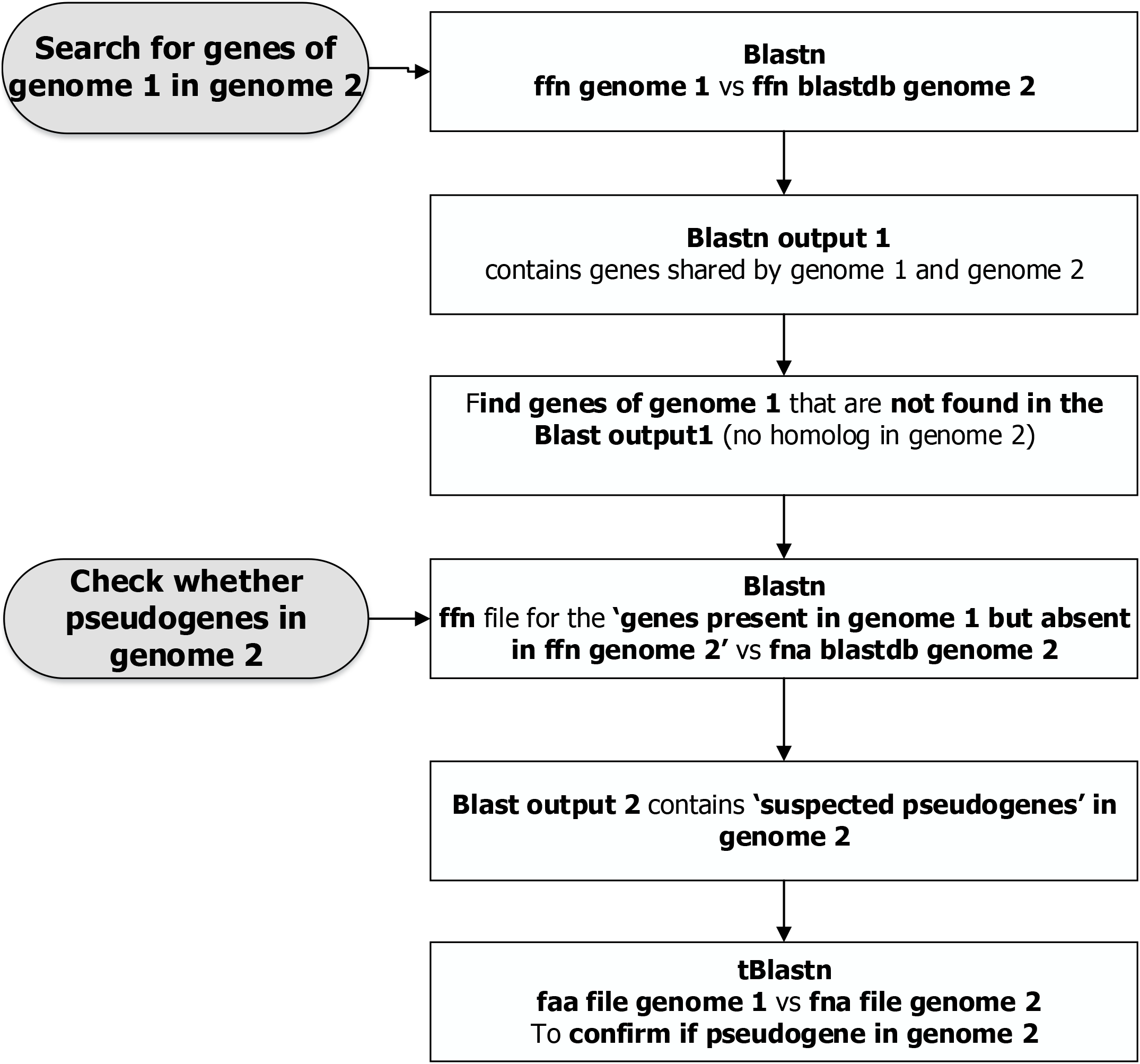
Methodology for finding gene presence-absence between two genomes.

The ffn files of H37Rv, the three L5 completed PacBio genomes and the L6 complete genomes, were also aligned using progressive Mauve (ver 20150226)(Darling et al. 2004) to identify rearrangements and examine synteny.

#### L5 Illumina (short reads) genomes

In total 355 L5 isolates genomes sequenced on the Illumina platform from various countries were included in the study. These genomes were derived from a larger study on genetic diversity of L5 and L6 (Coscolla et al. 2020). After reducing isolate redundancy (i.e. one representative retained for those that were extremely closely related), 205 L5 isolates genomes formed a non-redundant dataset. These 205 isolates (genomes) originated from West, South, East and Central Africa (Table S8), but primarily (n=155) from two regions within West Africa: the Western part of West Africa (including The Gambia, Sierra Leone, Ivory Coast, Liberia, Guinea and Mali) and the Eastern part of West Africa (including Ghana, Benin, and Nigeria) (Coscolla et al. 2020). In general, L5 and L6 are geographically restricted to West and Central-Africa, making this L5 genome selection representative of all the geographical regions affected.

### Mapping of L5 Illumina reads to H37Rv and L5 completed PacBio genomes

Raw reads (fastq files) of the 205 non-redundant Illumina genomes were mapped respectively to H37Rv and each of the 3 L5 completed PacBio genomes, using the MTBseq pipeline (Kohl et al. 2018). The depth mapping coverage of the samples in the clinical WGS data Illumina genomes against the reference genomes (percent read mapping, unambiguous coverage mean) was compared between the three L5 completed PacBio genomes and between each L5 completed PacBio genome and H37Rv. Mapping statistics parameters such as percent unambiguous total base, uncovered bases, SNPs, deletions, insertions, substitutions, percent genes mapped to reference, were also compared using each L5 completed PacBio genome or H37Rv as a reference.

#### Checking unique versus missing genes in Illumina L5 genomes, and SNPs in those confirmed as L5 specific

The genes found present or absent in L5 based on genome comparisons of the three L5 completed PacBio genomes with H37Rv were checked for their expected presence or absence in the L5 Illumina genomes. Using the Position Tables produced by the MTBseq pipeline, a gene was considered absent if 95% of its position in the genome had less than 8 reads covering them. From this data, a gene presence/absence matrix was generated for L5 Illumina genomes mapped to H37Rv and each of the three L5 completed PacBio genomes. Genes found to be L5-specific (present in completed PacBio and Illumina genomes) were also checked for SNPs in these genes using each of the L5 completed PacBio genomes as a reference. The script used for undertaking this analysis can be found at https://github.com/conmeehan/pathophy.

### Calculating the effect of reference genome selection on pairwise SNP distances

Transmission analysis of MTBC isolates often involves clustering isolates together based on specific SNP cut-offs (Meehan et al. 2019, 2018; Walker et al. 2013). We assessed whether the selection of reference genome (H37Rv, PcbL5Ben, PcbL5Gam or PcbL5Nig) changed the clustering rate of L5 isolates. For this we used the non-redundant set of L5 genomes from a previous study of L5 diversity (Coscolla et al. 2020), which included the 205 detailed above and a further 145 isolates that were closely related to at least one of these 205 isolates. Additionally, the TB-Profiler online SRA search tool (https://tbdr.lshtm.ac.uk/sra) was used to identify a further five isolates that were not in this dataset, resulting in 355 isolates included in this clustering study.

SNP alignments of the Illumina-sequenced genomes were created by first mapping to each of the four reference genomes (H37Rv and three PcbL5). The Amend function of MTBseq does this automatically for H37Rv, including masking of repetitive regions, accounting for 10% of the genome (Cole et al. 1998), and exclusion of columns with 95% ambiguous calls. To undertake this for the three L5 reference genomes, repetitive regions were first determined from the annotations of the genes. All genes whose description contained one of the following words was excluded: integrase, PE family, PE-PGRS, phage, transposase. SNP alignments were then created using MTBseq as was done for H37Rv. Pairwise distance matrices, one for each alignment based on each reference genome, were created using a python script which can be found at https://github.com/conmeehan/pathophy. Loose transmission clusters were created from these matrices at a cut-off of one, five and twelve SNPs, as previously described (Meehan et al. 2018). Clustering rates and the presence of a strain in a transmission cluster was then compared across the four reference genome mapping approaches.

### Determination of putative function of genes differentially present or absent in the L5 completed PacBio genomes

To find the (putative) function of genes present or absent only in L5 strains, and genes present in L5 strains but pseudogenes in H37Rv/L6/*M. bovis* or vice-versa, the fasta sequence of the genes were searched against the NR database of NCBI using BLASTX-2.8.1+ (Altschul et al. 1997) as well as against the Tuberculist database (http://tuberculist.epfl.ch), Mycobrowser (https://mycobrowser.epfl.ch/) (Kapopoulou, Lew, and Cole 2011) and literature. Furthermore, the gene function group/class was found using the COG database (Tatusov et al. 2000) and Mycobrowser.

### Determination of the sublineage of the L5 strains

The sublineage of the strains was determined by looking for sublineage defining SNP in the L5 genomes as previously described (Coscolla et al. 2020; Ates et al. 2018). The sublineage defining SNP were searched in the SNP data generated using the MTBseq pipeline with H37Rv as the reference genome.

## Results

### Comparative analysis of the completed L5 PacBio genomes and other lineages reveal differences in gene content

The number of genes, including paralogs, in the completed L5 PacBio genomes was: 4189 in the PcbL5Ben genome, 4162 in PcbL5Gam, and 4134 in PcbL5Nig versus 4126 in H37Rv, 4126 in the reference L6 genome and 4059 in the reference *M. bovis* genome. Hence, the maximum gene count difference was 55 genes among the three L5 isolates from this study, 63 genes among the human-adapted lineages (H37Rv, L5, L6), and 130 genes among the human- and animal adapted lineage (H37Rv, L5, L6 and *M. bovis*).

The visualization of the structure of the genomes showed that all three L5 genomes had a region (herein called region A) that was absent in H37Rv (gap between 1683331-1716665 on Fig.2). Furthermore, the Nigerian L5 completed PacBio genome (PcbL5Nig) was missing an additional region (herein called region B, between 1966666-2000000 in Fig.2) present in the Benin and Gambian L5 genomes (Fig.2) and also present in H37Rv.

**Figure 2.**
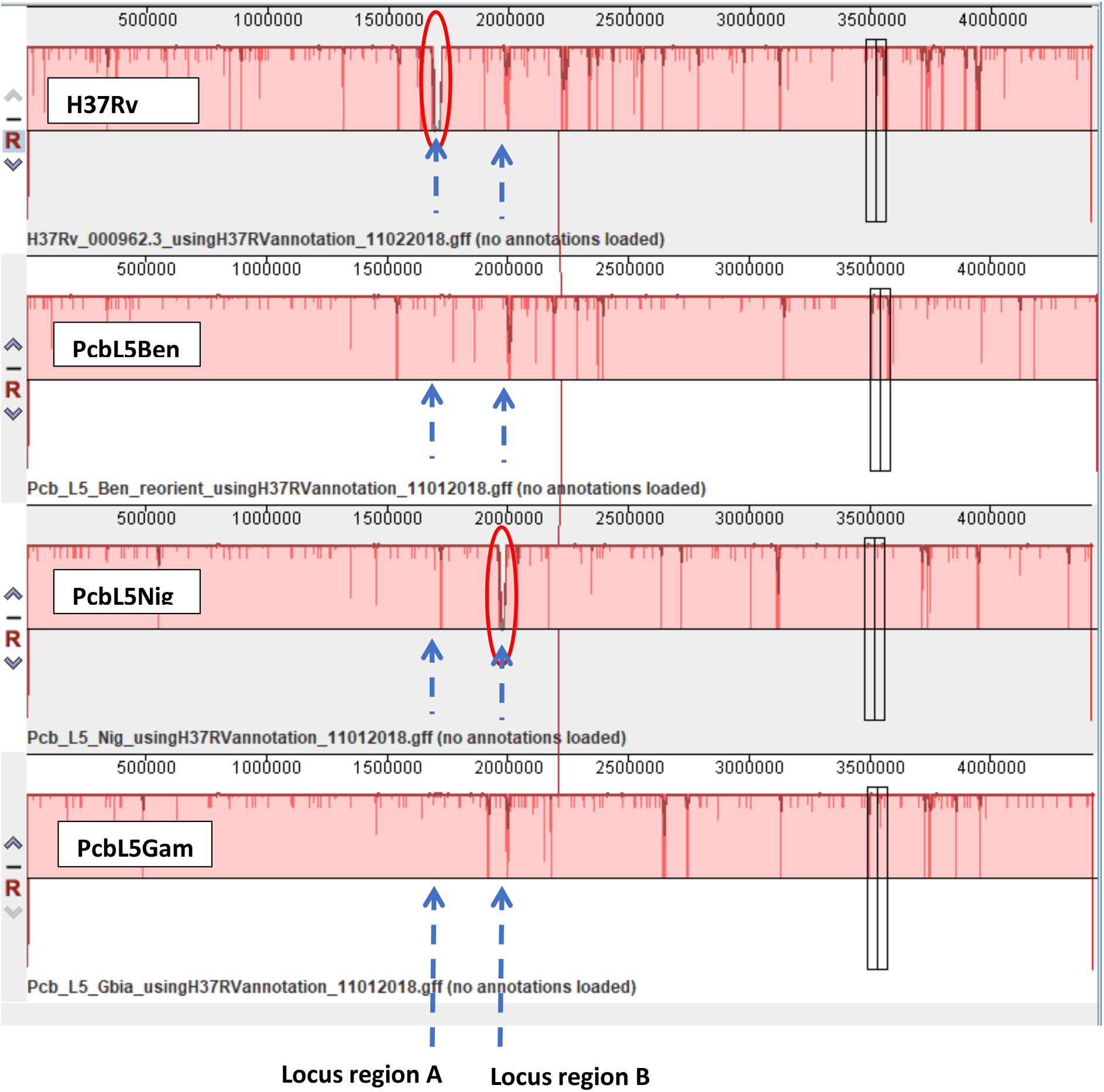
Visualization of possible rearrangements using Mauve pipeline: H37Rv (L4) genome and three completed PacBio genomes of *M. africanum* L5 isolates from Benin, The Gambia and Nigeria. The region circled in red (one on the H37Rv genome (missing region A), and one on the Nigerian completed PacBio genome (missing region B)) represents the regions absent in that strain genome but present in the other three strains complete genomes.

The Benin genome contained 5 genes (a block of 3 genes (1148 bp, 1104 bp, 2840 bp) and the other 2 (152 bp, 368 bp) randomly spread in the genome) that were neither present in the Nigerian or Gambian genomes, while the others contained respectively 1 (150 bp) and 2 (152 bp, 366 bp) unique genes randomly spread in the genome. These increased the pangenome of L5 by at least 6,280 bp. The Benin and Gambian strains each contained 33 genes that were not present in the Nigerian strain (Fig.3, Table 1, Table S1). Interestingly, 32 of these genes were also present in H37Rv (Fig.4, Table 1, Table S1). These 32 genes were thus missing in the Nigerian L5 strain exclusively, meaning that the default/ancestral state have the genes and then is loosing them. Those 32 genes-absent in the Nigerian L5 strain only-were sequentially contiguous and formed three blocks of 18, 12 and 2 genes (Table S1). We found that 30 of those genes, the blocks of 18 and 12 genes were separated by one gene and represented the region B (mentioned in the section above, Fig 2) found missing in the Nigerian L5 genome during another analysis (comparison of the structure of the 3 L5 completed PacBio genomes using Mauve pipeline). Ten genes (region A mentioned above) that were shared by the three L5 genomes were absent in H37Rv, while 9 genes (spread across the genome) present in H37Rv were not present in any of the three L5 genomes. Two genes (*Rv2073c*, *Rv2074*) of those 9 genes were only present in H37Rv but completely absent in the L5, L6 and *M. bovis* complete genomes.

**Figure 3.**
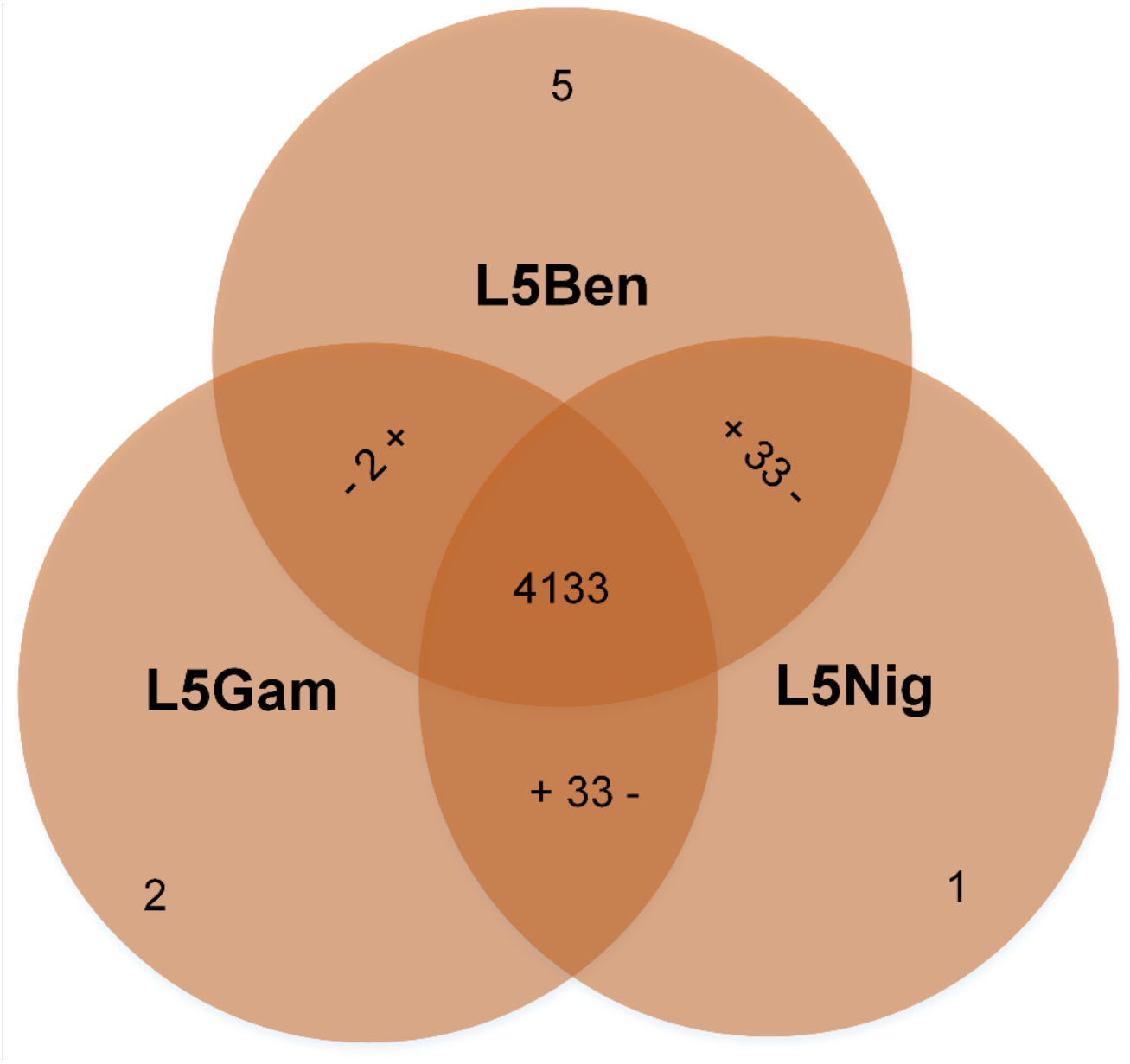
Gene count difference between three completed PacBio genomes of *M. africanum* L5 from Benin, The Gambia and Nigeria: L5Ben, L5Gam and L5Nig. The numbers are the count/number of gene difference (present or absent) between genomes. The sign (+) indicates the presence of genes while (−) indicates the absence of genes. For example the “− 2 +” gene difference between L5Gam and L5Ben, indicates that 2 genes are present in the L5Ben genome but absent in the L5Gam genome. 5, 2, 1 are the number of genes only found in respectively L5Ben, L5Gam, L5Nig.

**Table 1.** Gene content difference between *M. tuberculosis* H37Rv (L4) genome and completed PacBio genomes of three L5 isolates from Benin, The Gambia and Nigeria.

|  |  | Present in |  |  |  |
| --- | --- | --- | --- | --- | --- |
|  |  | PcbL5Ben | PcbL5Gam | PcbL5Nig | H37Rv |
| Absent in | PcbL5Ben |  | 2<sup>B</sup> | 1<sup>C</sup> | 9<sup>D</sup> |
|  | PcbL5Gam | 2 + 5<sup>A</sup> |  | 1<sup>C</sup> | 2 + 9<sup>D</sup> |
|  | PcbL5Nig | 33 + 5<sup>A</sup> | 33 + 2<sup>B</sup> |  | 32 + 9<sup>D</sup> |
|  | H37Rv | 10* + 5<sup>A</sup> | 10* + 2<sup>B</sup> | 10* + 1<sup>C</sup> |  |
|  |  | 4189 | 4162 | 4134 | 4126 |

**A:** include 5 only present in PcbL5Ben
**B:** includes 2 only present in PcbL5Gam
**C:** includes 1 only present in PcbL5Nig
**D:** includes 9 only present in H37Rv
**\*:** includes 10 genes shared by the PcbL5 (Ben, Gam, Nig) and absent in H37Rv

**Figure 4.**
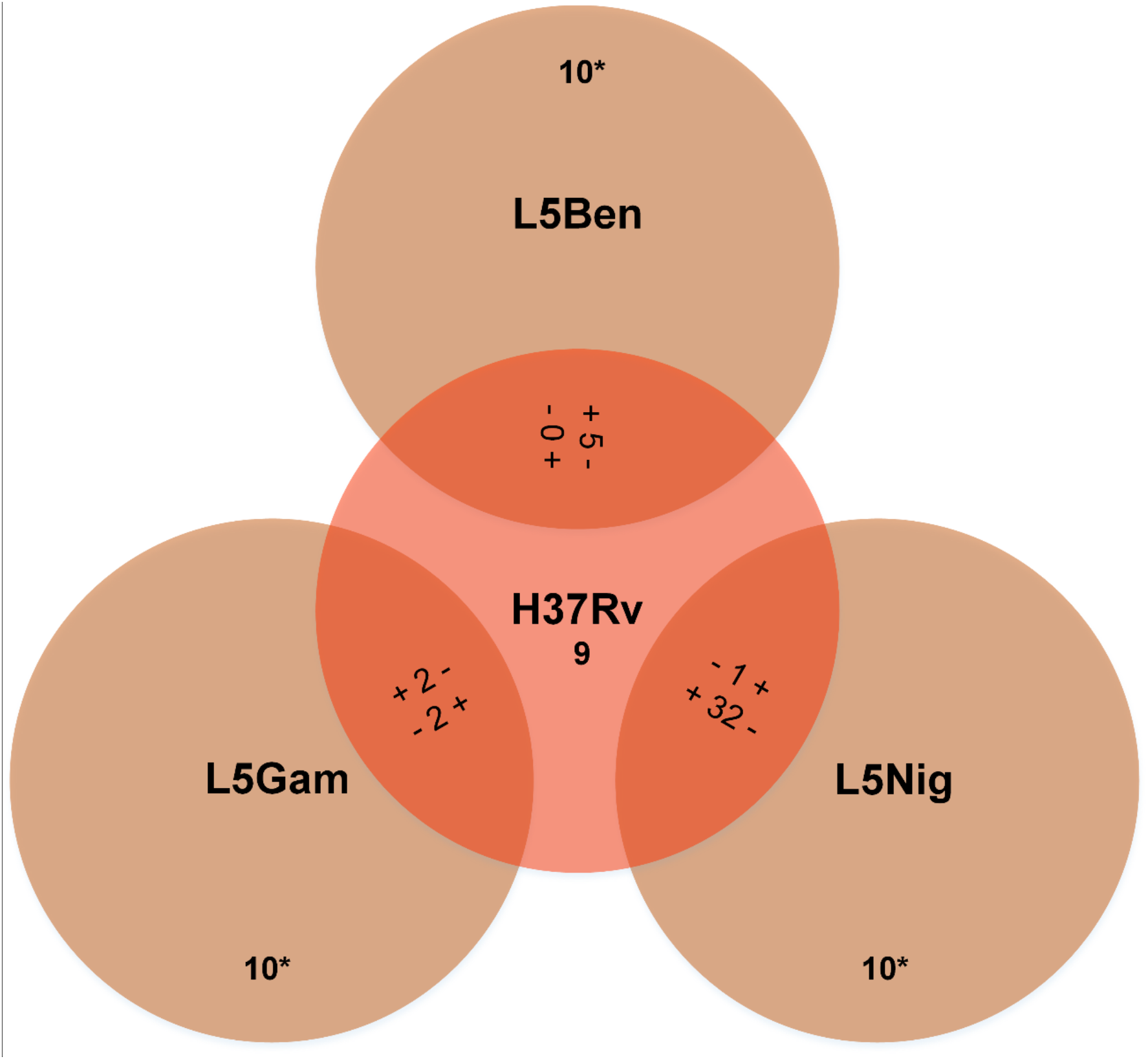
Gene count difference between *M. tuberculosis* H37Rv (L4) genome and three completed PacBio genomes of *M. africanum* L5 isolates from Benin, The Gambia and Nigeria. * These 10 genes are present in each of the three L5 completed PacBio genomes, and absent in H37RV.

Six of the genes shared by the three L5 completed PacBio genomes were confirmed pseudogenes in H37Rv (Table S2). Three genes present in H37Rv were confirmed to be pseudogenes in the three L5 completed PacBio genomes (Table S3).

### Gene presence/absence and related SNPs in lineage specific genes in a wider set of clinical isolates

The mapping estimates of the Illumina genomes are presented in Table 2. Mapping quality and coverage against a completed L5 reference was superior to the H37Rv reference approach (Table 2, Fig.5), as expected. Using the completed PacBio genomes from Benin and The Gambia had similar mapping estimates that were better than those of the Nigerian L5 genome, likely due to the large deletion in this genome. The genes specific to PcbL5Ben (5), PcbL5Gam (2), PcbL5Nig (1) were each found at various rates among the 205 L5 Illumina genomes (77.6%-97.1%, 159-199/205, Table 3).

**Table 2.** Mapping of 205 L5 isolates Illumina genomes to the *M. tuberculosis* H37Rv (L4) genome and completed genomes of three L5 isolates from Benin, The Gambia and Nigeria (mapping statistics/estimates). The best mapping results (numbers) are written in bold. When the best mapping result is obtained for PcbL5Nig as the reference, the next best result is also written in bold (as PcbL5Nig compared to the other two PcbL5 genomes missed a 30 genes region).

|  | H37Rv | PcbL5Ben | PcbL5Gam | PcbL5Nig |
| --- | --- | --- | --- | --- |
| Reads |  |  |  |  |
| Mean of Percent L5 reads mapped to | 96.9 | **97.5** | 97.3 | 96.5 |
| Mean of unambiguous coverage mean | 121.9 | 122.7 | **122.8** | 122.5 |
| Bases |  |  |  |  |
| Mean of Percent unambiguous total bases | 0.983 | 0.988 | **0.989** | 0.982 |
| Mean uncovered | 30444.8 | 18801.7 | **14657.5** | 35215.4 |
| Mean SNP | 2210 | 529.9 | **513.2** | **503.7** |
| Mean deletions | 373.3 | 95.9 | **95.7** | 97.2 |
| Mean insertions | 239.2 | **77.3** | 107.6 | **60.8** |
| Mean substitutions (including stop codons) | 1193 | **0** | **0** | **0** |
| Genes |  |  |  |  |
| Mean of Percent gene mapped (presence) | 99.59 | 99.72 | **99.79** | 99.76 |
| L5 illumina having all the ref. genome genes % (n=205) | 0 | 2 (4) | 7.3 (15) | 0 |
| Mean gene count difference between L5 Illumina genomes (gene count per Illumina-sequenced genome minus minimum gene count) | 29.8 | **38.3** | 33.4 | 33 |

**Figure 5.**
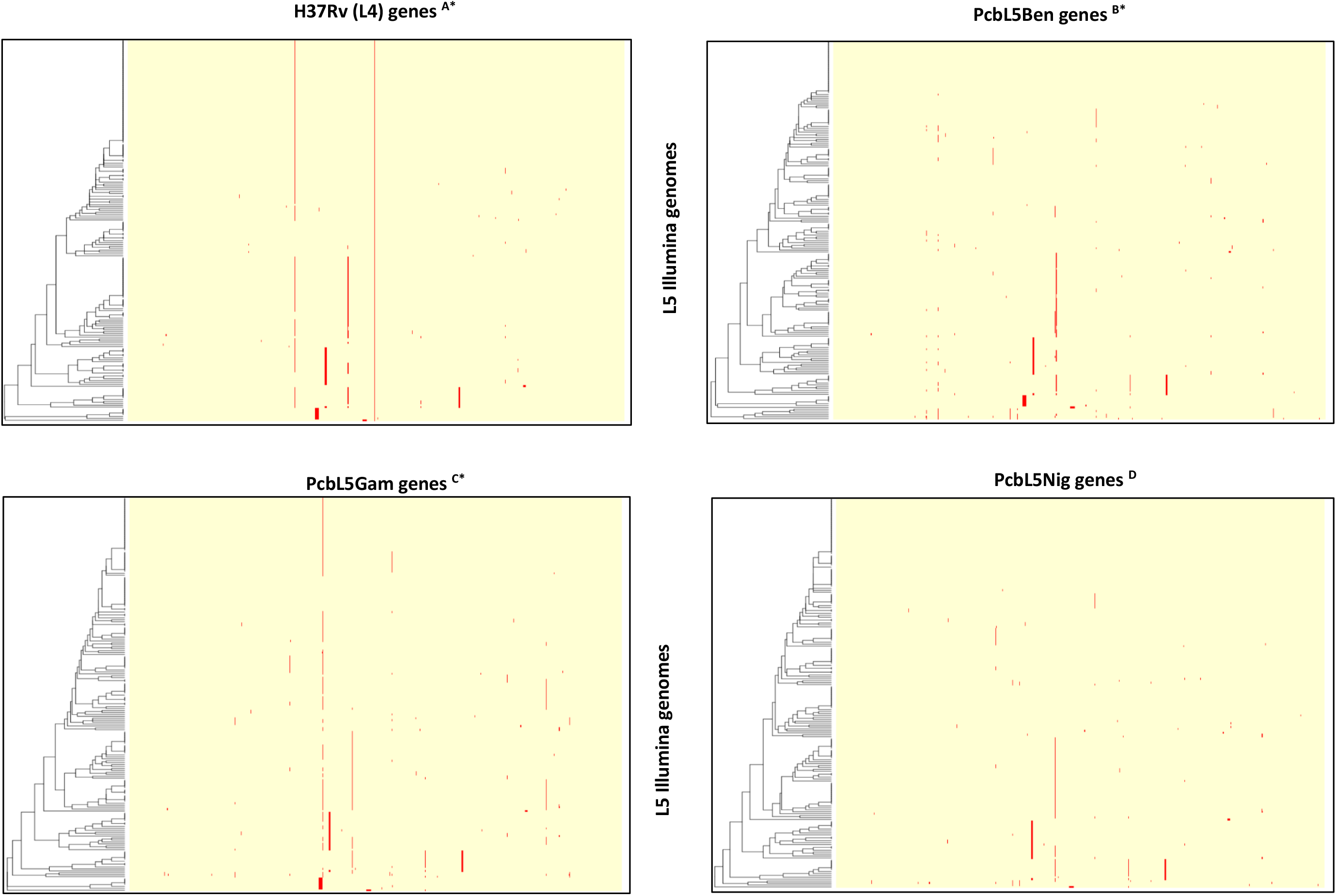
Mapping of 205 clinical (Illumina platform sequenced) genomes (L5_ig) of *M. africanum* L5 from various countries to *M. tuberculosis* H37Rv (L4) and three *M. africanum* L5 isolates (completed PacBio genomes) from Benin, The Gambia and Nigeria as reference (PcbL5Ben; PcbL5Gam; PcbL5Nig). (A) **5**/9 genes present in H37Rv but absent in the three L5_cg were **absent in all 205 L5_ig**. (B) **5**/5 genes **present in PcbL5Ben but absent in the two other L5 completed PacBio genomes** were **present in 159-199/205 L5_ig**. (C) **2**/2 genes **present in PcbL5Gam but absent in the two other L5 completed PacBio genomes** were **present in 195 & 199/205 L5_ig**. (D) **1**/1 gene **present in PcbL5Nig but absent in the two other L5 completed PacBio genomes** was **present in 195/205 L5_ig**. * The 32-33 genes absent in PcbL5Nig but present in PcbL5Ben, PcbL5Gam and *H37Rv* were also absent in 6/205 L5_ig (Fig.9). **…. or ….** gene absent

**Table 3.**
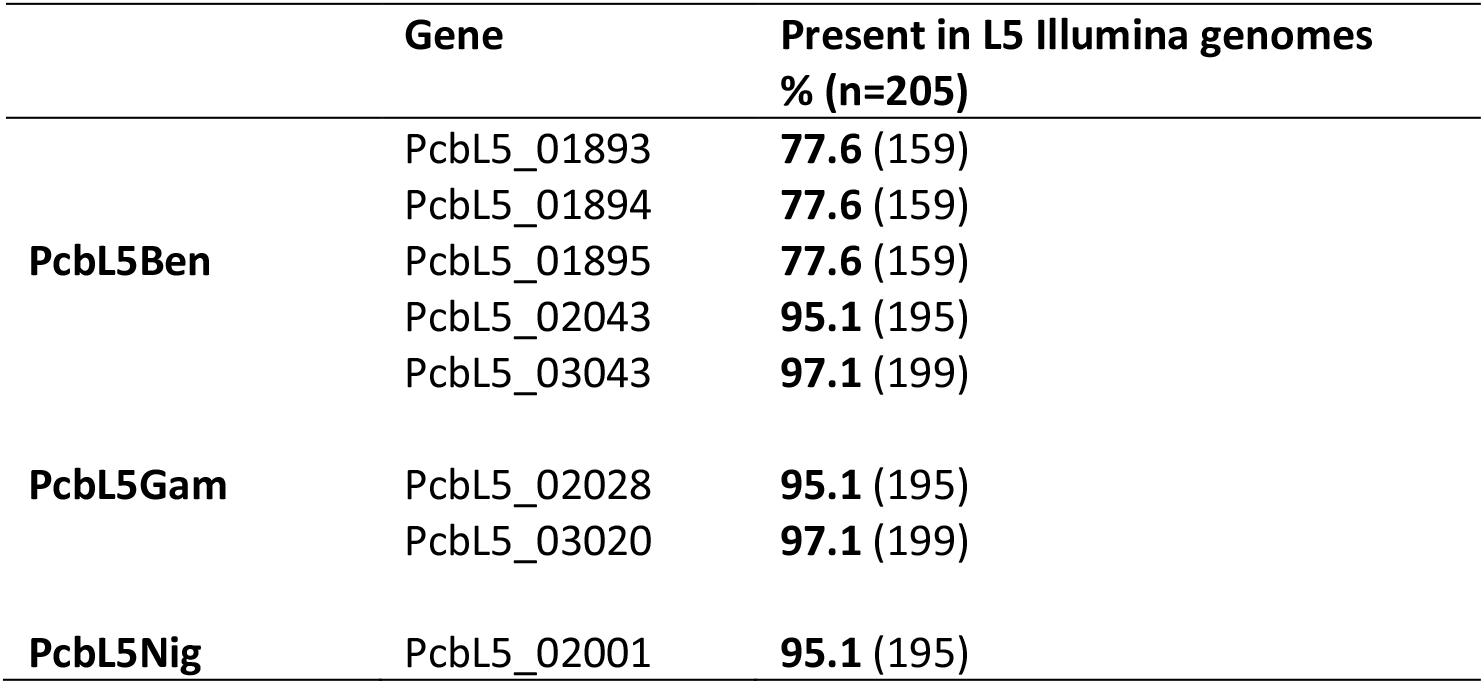
Presence in 205 L5 isolates Illumina genomes of genes detected in only one of the three L5 completed genomes from Benin, Nigeria or The Gambia.

Interestingly, 2.9% of the Illumina L5 genomes (6/205) had similar patterns of large gene loss as the Nigerian completed PacBio genome as they missed 30 of the 33 genes present in the Benin and Gambian genomes and H37Rv. These six PcbL5Nig-like Illumina L5 genomes formed a monophyletic group, within the L5 clade (Fig. 6), suggesting a single loss of these gene clusters, although those six L5 genomes originated from several different countries including Benin, Ghana and Nigeria. The two blocks of 18 (14835 bp, *Rv1493* through *Rv1509*) and 12 genes (10209 bp, *Rv1511* through *Rv1521*), separated by one gene (3441 bp, *Rv1510*) amounted in total to 25044 bp which is around 0.6% of the length (around 4 million bp) of a typical MTBC (H37Rv) genome. The two blocks contain genes whose annotations include *mutB*, *mazE4*, *mazF4* and others with various putative functions (Table S4).

**Figure 6.**
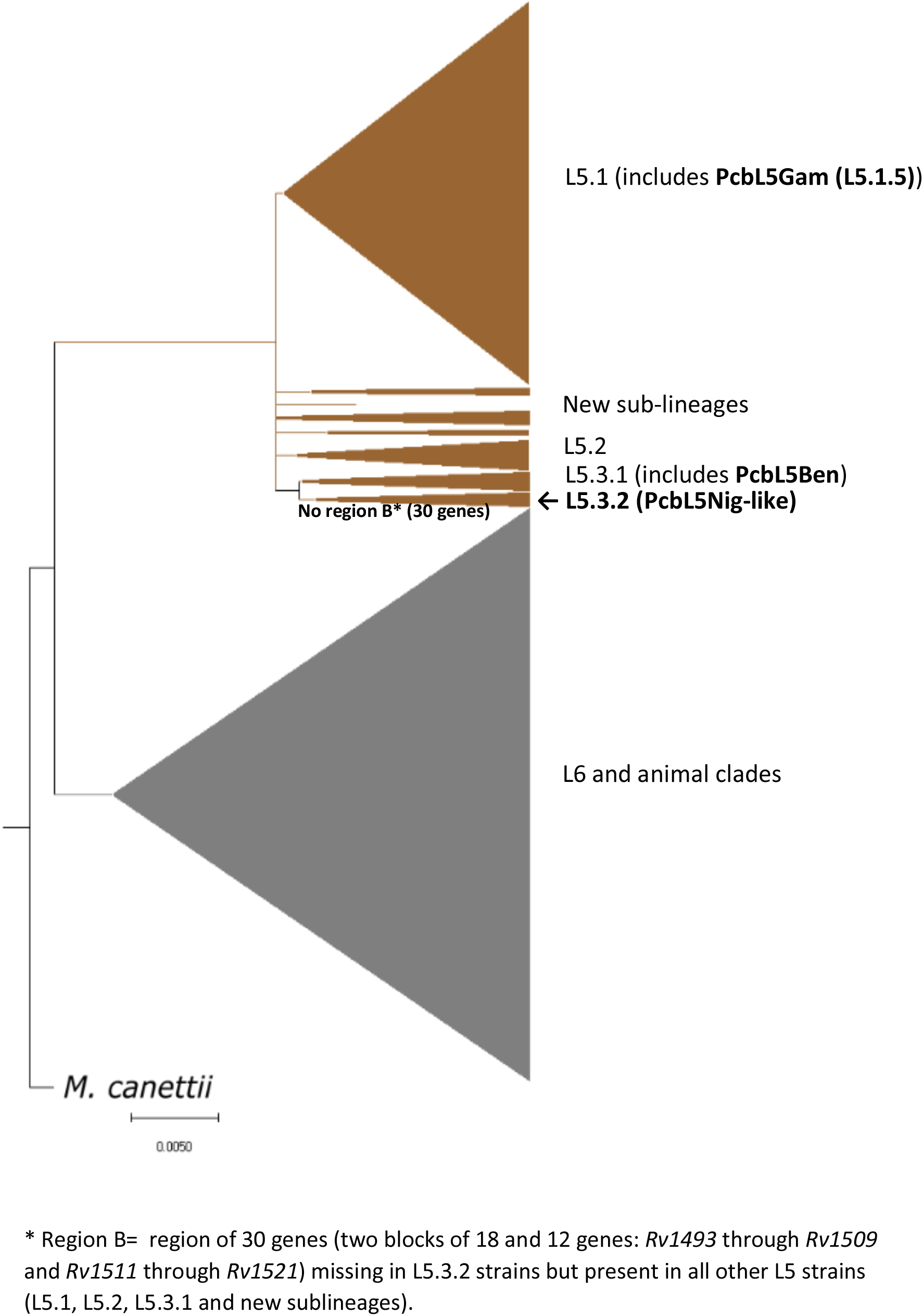
Phylogenetic tree showing the 6 L5 isolates Illumina genomes (L5.3.2) similar to the completed PacBio genome of the Nigerian L5 strain (PcbL5Nig) and the position of the other two L5 PacBio genomes (PcbL5Ben and PcbL5Gam)

Four of the ten genes present in the three completed L5 PacBio genomes but absent in H37Rv were found in all the 205 L5 Illumina genomes while the other were found in variable amounts 81-98% (165-201) (Table S5). These four gene amounted to a total of 2382 bp (Table S5), representing around 0.05% of the length of a typical MTBC genome. These four genes include: *Mb2048c*, PE35, a hypothetical protein-possibly an IS*256* transposase, and a hypothetical protein possibly related to the CAAX conserved domain. Two of these four genes were present in L5 genomes only (PE35 and hypothetical protein possibly CAAX conserved domain) while the remaining two (*Mb2048c* and hypothetical protein possibly IS256 transposase) were found in L6 and *M. bovis* as well (Table S5). Importantly for phylogenetic purposes, SNPs were detected in all four genes in 1.5-3.4% of the 205 L5 Illumina genomes (Table S5). Predicted functions for all L5-specific genes are listed in table S5.

Five (*Rv1977*, *Rv1979c*, *Rv1993c*, *Rv1995*, *Rv2073c*) of the 9 genes present in H37Rv and absent in the three completed L5 PacBio genomes were absent in all the 205 L5 Illumina genomes. These five genes amounted to a total of 4,245 bp (Table S6), representing around 0.1% of the length of a typical MTBC genome. Four of these genes (*Rv1977*, *Rv1979c*, *Rv1993c*, *Rv1995*) were absent in L5 strains only (i.e. present in L6 and *M. bovis* reference genomes) while the fifth (*Rv2073c*) was absent in L6 and *M. bovis* as well (Table S6). The four other genes absent from the completed PacBio genomes were present in a minority of the L5 Illumina genomes (Table S6, Fig. 5). Predicted functions for all nine genes are listed in table S6.

### Sublineage of the strains (PacBio completed and Illumina genomes)

The determination of the strains sublineage revealed that the PcbL5Ben is an L5.3 strain that contains the 30 genes (two blocks of 18 and 12 genes: *Rv1493* through *Rv1509* and *Rv1511* through *Rv1521*) whereas PcbL5Nig is also a L5.3 strain, but situated in a different clade (Fig. 6). Based on this region (a new RD), we classified PcbL5Ben as an L5.3.1 strain and the PcbL5Nig as a L5.3.2 strain; the PcbL5Gam isolate is an L5.1.5 strain. The Illumina genomes were distributed as follows (Table S7): 90 L5.1.1; 17 L5.1.2; 12 L5.1.3; 21 L5.1.4; 9 L5.1.5 amounting to a total of 149 L5.1 strains (72.7%); 13 (6.3%) L5.2 strains; 8 L5.3.1; 6 L5.3.2 amounting to a total of 14 L5.3 strains (6.8%) and 29 (14.2%) strains of unknown sublineages (potentially new sublineages). All L5 strains including L5.1, L5.2, L5.3.1, unknown sublineages, except L5.3.2 strains have that 30 genes region.

### Changes to transmission clustering rates based on selection of reference genome

Transmission analysis was undertaken on an expanded set of 355 L5 isolates using each of the four reference genomes (H37Rv, PcbL5Ben, PcbL5Gam and PcbL5Nig) for the SNP mapping. The current gold standard is to use H37Rv for calling SNPs and then creating transmission clusters at a specific SNP cut-off. For this dataset, using a 12 SNP cut-off, it was found that 40.6% (n=144) isolates were within a transmission cluster with at least one other isolate (Table 4), for a otal of 55 clusters. Substituting PcbL5Ben instead of H37Rv as the reference genome reduced the number of clusters to 54, removing 3 isolates from transmission clusters, resulting in a clustering rate of 39.7% at a 12 SNP cut-off (Table 4). A similar reduction in transmission clustering was observed when using the PcbL5Nig genome as a reference but not when using the PcbL5Gam. When a more conservative SNP cut-off was used, such as 1 SNP or 5 SNPs, the reduction in transmission clustering rates was more pronounced and was observed for all the PcbL5 genomes (Table 4). For example, at a 1 SNP cut-off, H37Rv-based mapping resulted in 100 isolates (28.2%) being placed inside transmission clusters, but this reduced to 94 or 95 isolates (26.5-26.8%) when using a L5 genome as the reference. This demonstrates that even on this small dataset, transmission cluster estimations are affected by the selection of reference genome, with potential over-estimation of recent transmission when using H37Rv as the reference genome.

**Table 4.** Comparison of transmission clustering rates based on choice of reference genome. Short read data from 355 L5 isolates were mapped against each of the four reference genomes for SNP calling. Distance matrices between all isolates were constructed per reference approach and transmission clusters were defined based on specific SNP cut-offs

1 SNP
| Reference used | In transmission cluster | Percentage of total dataset | Number of clusters |
| --- | --- | --- | --- |
| H37rv | 100 | 28.17 | 44 |
| PcbL5Ben | 94 | 26.48 | 43 |
| PcbL5Gam | 95 | 26.76 | 43 |
| PcbL5Nig | 95 | 26.76 | 43 |

5 SNP
| Reference used | In transmission cluster | Percentage of total dataset | Number of clusters |
| --- | --- | --- | --- |
| H37rv | 129 | 36.34 | 53 |
| PcbL5Ben | 124 | 34.93 | 51 |
| PcbL5Gam | 124 | 34.93 | 51 |
| PcbL5Nig | 124 | 34.93 | 51 |

12 SNP
| Reference used | In transmission cluster | Percentage of total dataset | Number of clusters |
| --- | --- | --- | --- |
| H37rv | 144 | 40.56 | 55 |
| PcbL5Ben | 141 | 39.72 | 54 |
| PcbL5Gam | 144 | 40.56 | 55 |
| PcbL5Nig | 141 | 39.72 | 54 |

## Discussion

Comparison of completed PacBio and Illumina short-reads genomes from clinical isolates revealed gene content diversity within L5 isolates as well as between L5 and other lineages.

We found that four genes, present in all the L5 isolates of our multi-country collection, were absent in the H37Rv genome. Those genes represent 2,382 bp which is around 0.05% of the total length (base-pair) of a typical MTBC genome. Furthermore, those genes contain SNPs that can be used to identify additional diversity between L5 strains. This has implications when using precise SNP cut-offs to define transmission clusters (Walker et al. 2013; Meehan et al. 2018) as SNPs in these genes would be missed with H37Rv mapping, resulting in underestimation of genome diversity of L5 strains. Our findings suggest, that using an L5 genome as a reference would likely increase the resolution of L5 genome analyses over the H37Rv approach and can potentially increase the discrimination power, leading to a reduction of the amount of recent L5 transmission inferred.

Our findings are in line with other reports indicating that some genes present in genomes of clinical MTBC strains were absent in the H37Rv genome (Periwal et al. 2015; O’Toole and Gautam 2017), and with other recommendations to use additional reference genomes different to H37Rv (Norman et al. 2019; O’Toole and Gautam 2017; Lee et al. 2020). Although Lee *et al*. (Lee and Behr 2016) - after comparing various lineages as reference genome for L4 genome analysis - concluded that there is no need to use a lineage-specific reference genome, their observation was only based on the analysis of L4 clinical isolates and focused on SNPs and short indels, not larger gene deletions or SNPs within these regions. In contrast, our findings indicate that mapping of NGS data from L5 strains to L4 reference will have an impact, in terms of both reference genome coverage and coverage of lineage-specific genes.

Although the use of an L5 genome as reference would have many benefits over H37Rv for particular study questions, several gene content differences were still observed within the L5 lineage. Each of the three L5 PacBio genomes had genes unshared with the two others that were all found in L5 Illumina genomes (thus excluding exogenous contamination). The absence of 30 genes in the Nigerian L5 completed PacBio genome (but shared by the Benin and Gambian long read genomes), also found in six L5 genomes from different countries, confirmed the large deletions in the PcbL5Nig genome and showed that it is representative of some L5 strains in some countries. Deletions in the genomes of strains of MTBC lineages were previously reported (Tsolaki et al. 2005; Gagneux et al. 2006; Kato-Maeda et al. 2001). The fact that, on the phylogenetic tree the six isolates formed a monophyletic group of the (SNP defined) L5.3 sub-lineage (Coscolla et al. 2020) suggests that the loss of these 30 genes (*Rv1493* through *Rv1509*, and *Rv1511* through *Rv1521*) is a marker of the L5.3.2 sub-lineage, compared to the sublineage L5.3.1 strains that have these genes intact. Thus, we find that regions of difference (RDs) are present both between lineages and between sub-lineages.

We found that the genes *Rv2073c* and *Rv2074* that were absent in L5, L6 and *M. bovis*. It was previously reported that L5, L6 and *M. bovis* all lack the RD9 region (Brosch et al. 2002; Mostowy et al. 2002; Gagneux et al. 2006), which contains *Rv2073c*, *Rv2074* and partly *Rv2075* (Gordon et al. 1999). The lack of RD9 by L5, L6 and *M. bovis* is thus confirmed by our data, however one (1/205) of the L5 genomes had *Rv2074* present. Of the five genes (*Rv1977*, *Rv1979c*, *Rv1993c*, *Rv1995*, *Rv2073c*) absent in all L5 strains, besides *Rv2073c* included in RD9, one (*Rv1977*) was part of the RD7 region (Gordon et al. 1999), another (*Rv1979c*) was part of the RD2 region (Gordon et al. 1999), and *Rv1993c* and *Rv1995* potentially included in other RD regions. These findings show that gene presence/absence approach sheds more light on the differences between lineages than the RD approach alone would do.

The classification of the L5-specific genes into functional groups based on Mycobrowser and Tuberculist databases (Table 4) revealed a variety of functions including: a PE/PPE protein family (possibly PE35) which may be linked to antigenic variation (NCBI annotation) and virulence (STRING database annotation (Szklarczyk et al. 2017)); two ABC-transporters; an IS*256*-like transposase, whose homology (IS256) is a marker of specific virulent lineages of staphylococci (Murugesan et al. 2018; Gu et al. 2005); and a gene with a CAAX conserved domain, implicated in post-translation modification or signal transduction (Gao, Liao, and Yang 2009a, 2009b). Only one (*Rv1979c*) of the four genes absent in only L5 strains had a predicted function, labelled as “cell wall and cell processes” (Table S6). While *Rv1979c* is associated with clofazimine and bedaquiline resistance (Zhang et al. 2015; Ghodousi et al. 2019), two of the drugs used for the treatment of RIF-resistant/MDR-TB (WHO 2016, 2019), minimal inhibitory concentration testing of clofazimine in five L5 strains did not find the deletion conferred resistance (Merker et al. 2020). Further studies and protein function discovery is needed to investigate the consequences of the absence of those genes in L5.

The genes *Rv1978* and *Rv1994c* present in H37Rv and the completed reference genomes of L6 and *M. bovis*, were absent in the 3 completed Pacbio genomes of L5 strains, yet present in few L5 isolates Illumina genomes (3/205, 1.5% for *Rv1978*; 1/205, 0.5% for *Rv1994c*). *Rv1978* is required for bacterial survival in macrophages, and non-essential for *in vitro* growth of H37Rv (Rengarajan, Bloom, and Rubin 2005). *Rv1994c* is involved in the regulation and transport (efflux) of toxic metals, especially copper, which is toxic in excess (Samanovic et al. 2012) and may hamper *in vitro* growth, and survival during the chronic phase of infection, similar to the effect of disruption of *csoR* (Marcus et al. 2016; Rowland and Niederweis 2012; Ward, Hoye, and Talaat 2008; Cavet et al. 2003) (Table S6).

The functional annotation for the H37Rv-specific genes *Rv2073c* (short chain dehydrogenase) and *Rv2074* (pyridoxamine-5-phosphate oxydase: vitamin B6 (pyridoxine)) absent in L5, L6 and *M. bovis* strains is “intermediary metabolism and respiration”. Those two genes are part of the RD9 region, which confirmed their absence in L5, L6 and *M. bovis* that have that region deleted (Gordon et al. 1999). *Rv2073c* is an NAD(P)-dependent oxidoreductase (NCBI) that catalyzes a wide range of reactions and is involved in redox sensor mechanisms (Kavanagh et al. 2008; Sellés Vidal et al. 2018). *Rv2074* was previously thought to be a pyridoxine (vitamin B6) oxidase, but is now known to be a F420 dependent biliverdin reductase, a cofactor of vitamin B6 synthesis (UniProt) (Ahmed et al. 2016; Selengut and Haft 2010). Vitamin B6 is essential for survival and virulence of *M. tuberculosis* (Dick et al. 2010) and its cofactor (F420 dependent biliverdin reductase) is implicated in immune-evasive mechanisms to allow bacterial persistence (Ahmed et al. 2016; Selengut and Haft 2010)(Table S6). Recently, it has been reported that the synthesis of F420 might be stimulated by phosphoenolpyruvate (Grinter et al. 2020), which is the precursor to pyruvate, a supplement often added to culture medium to improve the *in vitro* growth of L5, L6 and *M. bovis*. Furthermore, there is a suggestion that F420 is needed for the activation of the antituberculosis drugs pretomanid and delamanid (Grinter et al. 2020).

The absence of these genes in most L5 isolates suggests that L5 strains would be less likely to survive in macrophages (*Rv1978*), have reduced growth *in vitro* (*Rv1994c*, as previously found by Sanoussi et al (Sanoussi et al. 2017)), be less immune-evasive, less persistent and less virulent (*Rv2074*) than L4 strains (at least H37Rv; as previously suggested for L5 and L6 strains (Coscolla and Gagneux 2014; Reiling et al. 2013)). In addition, the absence of *Rv2074* in most L5 (and in the complete genomes of L6 and *M. bovis* as well) suggests that L6 and *M. bovis* strains would also be less likely to be immune-evasive, to be persistent and to survive than their L4 counterparts. Despite the presence of vitamin B6 in boiled egg (thus Lowenstein-Jensen medium), its quantity in the L.J medium is probably insufficient to allow a high yield of L5, L6 and *M. bovis* growth in culture (survival). Further studies, including vitamin B6 supplementation, could investigate the consequences of the absence of those genes in most L5 strains (and L6 and *M. bovis* for *Rv2074*).

Transmission clustering analyses revealed differences in clustering rates and inclusion of isolates in clusters based on the selection of the reference genome (Table 4). Overall, H37Rv-based mapping was found to place a small percentage more isolates in transmission clusters than any L5-based mapping approach. This is due to the additional genes found in these genomes also being present in the clinical isolates and these genes containing differentiating SNPs. Thus, with an increased discovery of SNPs in non-H37Rv genes, some isolates that previously satisfied the SNP distance criterion for clustering were now excluded as they are less similar than they appear to be when using H37Rv as a reference. This was most pronounced at the lower SNP cut-offs (Table 4). This has strong implications for molecular epidemiology in West Africa, where most L5 strains are found. Additionally, if similar scenarios exist for other lineages, this may result in changes to all non-L4 transmission analyses.

The within-lineage differences in both gene content and potential functionality indicates a need for more closed genomes of MTBC sub-lineages and that perhaps a sub-lineage-specific reference genome approach would be appropriate for MTBC WGS and molecular epidemiology. However, although an *ad hoc* (e.g. outbreak-specific) reference genome provides a more accurate and precise comparison of strains involved in a specific situation, that reference is only specific to that particular situation/outbreak/population/lineage and cannot be used in another context, making comparisons between lineages and settings difficult.

As an alternative to using a single MTBC-wide, lineage or sub-lineage specific reference genome approach, the variable presence of genes within one lineage suggests that an L5 pangenome-based reference genome capturing all the known diversity may be required instead. Such a genome would be a composite of all the genes found in all lineages and sub-lineages of the MTBC (i.e. both the core and accessory genome) (McInerney et al. 2020) and is usually represented as a graph instead of a single sequence (Rakocevic et al. 2019; Paten et al. 2017; Martiniano et al. 2019; Marschall et al. 2018; Church et al. 2015). Alternatively, it can be a selection of reference genomes, with mapping to all or a subset undertaken, as has been done with *M. chimaera* and other pathogens (van Ingen et al. 2017; Garimella et al. 2019). Other authors have also reported that because of the genetic variability between strains, using a single strain genome as reference genome lacks accuracy (Ballouz, Dobin, and Gillis 2019; Sherman et al. 2019; Li et al. 2010). This MTBC-wide pangenome approach has been suggested before, including for other organisms (Periwal et al. 2015; Meehan et al. 2019; Jandrasits et al. 2019; Ballouz, Dobin, and Gillis 2019; Medini et al. 2005; Tettelin et al. 2005; Sherman et al. 2019; Li et al. 2010). However, such an approach also has its own drawbacks, including difficulty in mapping reads that bridge the boundaries between accessory genes and the rest of the genome, comparing strains of different lineages including phylogenetic analyses and retention of gene names and codes in clinical use, where H37Rv is deeply embedded (Marschall et al. 2018). The specific sequence of each gene would also need to be chosen for such a reference, with the inferred ancestral genome representative of MTBC lineages approach being the most likely method (Goig et al. 2020, 2018; Comas et al. 2013)

Another alternative approach is a *de novo* assembly reference free approach (Maretty et al. 2017; Iqbal et al. 2012; Meehan et al. 2019; Li et al. 2010) where strains are either assembled into contigs without the use of a reference genome, or compared to each other without first calling SNPs. This would allow for clustering of samples regardless of lineage, e.g. for genotyping or transmission analyses using Mash (software for fast/meta -genome distance estimation technique) (Ondov et al. 2016) but would require new cut-offs for defining clusters and many additional steps for further gene annotation and between sample comparisons, making its clinical use potentially confusing.

A limitation of this study is that the completed (PacBio) and short-read (Illumina) genomes were derived from positive cultures, excluding possible minority L5 strain diversity as L5 strains are overrepresented in negative cultures (Sanoussi et al. 2017). WGS applied directly on sputum is more and more needed, especially for ancestral lineages (including L5 and L6) where growth in culture is problematic (negative culture or dysgonic isolates that are a challenge to grow for DNA extraction (Sanoussi et al. 2018, 2017)). Also, a limitation regarding the comparison of gene presence/absence in L5 versus L4 (including H37Rv), L6, *M. bovis* complete genomes, is that our study included only a single complete genome of L4, L6 and *M. bovis*, while these lineages may also display variability similar to the intra-L5 variability we observed in our study.

In conclusion, the use of a (sub-)lineage reference genome can increase the resolution for strains comparison in comparison to a H37Rv based mapping approach for L5 genome analyses for epidemiology (transmission), phylogeny and sub-lineage determination. Still, the use of a (sub-)lineage reference genome may miss some within-lineage gene differences. For drug-resistance detection in clinical L5 strains or strains of other lineages, H37Rv could still be used as reference genome as resistance-related mutations are usually among core genes (shared across all lineages). The high within-lineage gene content variability suggests the pangenome of MTBC strains may be larger (at least by 6,280 bp) than previously thought, implying a reference-free genome assembly (*de novo assembly*) approach may be needed.

## Authors and contributors

CNS was involved in conceptualization, methodology, formal analysis, investigation, data curation, writing of the original draft, review/editing the manuscript and visualization.

MC was involved in conceptualization, methodology, investigation, data curation, review and editing the manuscript and visualization.

BO was involved in methodology, investigation and review/editing the manuscript.

IDO was involved in resources and review/editing the manuscript

MA was involved in resources.

SN was involved in conceptualization, reviewing/editing the manuscript and supervision.

JP was involved in investigation, data curation and reviewing/editing the manuscript.

SH was involved in conceptualization, investigation, data curation and reviewing/editing the manuscript.

DY was involved in conceptualization, resources and reviewing/editing the manuscript.

SG was involved in conceptualization and reviewing/editing the manuscript.

LR was involved in conceptualization, resources, review/editing the manuscript and supervision.

DA was involved in resources, review/editing the manuscript and supervision.

BCdJ was involved in conceptualization, methodology, writing of the original draft, review/editing the manuscript and supervision.

CJM was involved in conceptualization, methodology, formal analysis, investigation, writing of the original draft, review/editing the manuscript, supervision and visualization.

## Ethical approval

The PacBio sequenced L5 strain from Benin was isolated during the multicentric OFLOTUB study approved by the National Ethics Committee in Cotonou, Benin and each of the participating countries (Merle et al. 2012).

## Acknowledgements

We would like to acknowledge input from Pim de Rijk Willem and Patrick Beckert for their technical inputs.

## Reference

Ahmed, F Hafna, A Elaaf Mohamed, Paul D Carr, Brendon M Lee, Karmen Condic-Jurkic, Megan L O’mara, and Colin J Jackson. 2016. “Rv2074 Is a Novel F 420 H 2-Dependent Biliverdin Reductase in Mycobacterium Tuberculosis.” https://doi.org/10.1002/pro.2975.

Altschul, Stephen F., Warren Gish, Webb Miller, Eugene W. Myers, and David J. Lipman. 1990. “Basic Local Alignment Search Tool.” Journal of Molecular Biology 215 (3): 403–10. https://doi.org/10.1016/S0022-2836(05)80360-2.

Altschul, Stephen F, Thomas L Madden, Alejandro A Schäffer, Jinghui Zhang, Zheng Zhang, Webb Miller, and David J Lipman. 1997. “Gapped BLAST and PSI-BLAST: A New Generation of Protein Database Search Programs.” Nucleic Acids Research. Vol. 25. Oxford University Press. https://www.ncbi.nlm.nih.gov/pmc/articles/PMC146917/pdf/253389.pdf.

Ates, Louis S, Anzaan Dippenaar, Fadel Sayes, Alexandre Pawlik, Christiane Bouchier, Laurence Ma, Robin M Warren, et al. 2018. “Unexpected Genomic and Phenotypic Diversity of Mycobacterium Africanum Lineage 5 Affects Drug Resistance, Protein Secretion, and Immunogenicity.” Genome Biology and Evolution 10 (8): 1858–74. https://doi.org/10.1093/gbe/evy145.

Ballouz, Sara, Alexander Dobin, and Jesse A. Gillis. 2019. “Is It Time to Change the Reference Genome?” Genome Biology. BioMed Central Ltd. https://doi.org/10.1186/s13059-019-1774-4.

Behr, M. A., M. A. Wilson, W. P. Gill, H. Salamon, G. K. Schoolnik, S. Rane, and P. M. Small. 1999. “Comparative Genomics of BCG Vaccines by Whole-Genome DNA Microarray.” Science 284 (5419): 1520–23. https://doi.org/10.1126/science.284.5419.1520.

Belisle, John T., and Michael G. Sonnenberg. 1998. “Isolation of Genomic DNA from Mycobacteria.” In Mycobacteria Protocols, 31–44. New Jersey: Humana Press. https://doi.org/10.1385/0-89603-471-2:31.

Bentley, Stephen D., Iñaki Comas, Josephine M. Bryant, Danielle Walker, Noel H. Smith, Simon R. Harris, Scott Thurston, et al. 2012. “The Genome of Mycobacterium Africanum West African 2 Reveals a Lineage-Specific Locus and Genome Erosion Common to the M. Tuberculosis Complex.” Edited by Pamela L. C. Small. PLoS Neglected Tropical Diseases 6 (2): e1552. https://doi.org/10.1371/journal.pntd.0001552.

Bifani, P, S Moghazeh, B Shopsin, J Driscoll, A Ravikovitch, and B N Kreiswirth. 2000. “Molecular Characterization of Mycobacterium Tuberculosis H37Rv/Ra Variants: Distinguishing the Mycobacterial Laboratory Strain.” Journal of Clinical Microbiology 38 (9): 3200–3204. https://www.ncbi.nlm.nih.gov/pubmed/10970357.

Boritsch, Eva C., Varun Khanna, Alexandre Pawlik, Nadine Honoré, Victor H. Navas, Laurence Ma, Christiane Bouchier, et al. 2016. “Key Experimental Evidence of Chromosomal DNA Transfer among Selected Tuberculosiscausing Mycobacteria.” Proceedings of the National Academy of Sciences of the United States of America 113 (35): 9876–81. https://doi.org/10.1073/pnas.1604921113.

Brites, Daniela, Chloé Loiseau, Fabrizio Menardo, Sonia Borrell, Maria Beatrice Boniotti, Robin Warren, Anzaan Dippenaar, et al. 2018. “A New Phylogenetic Framework for the Animal-Adapted Mycobacterium Tuberculosis Complex.” Frontiers in Microbiology 9 (NOV). https://doi.org/10.3389/fmicb.2018.02820.

Brosch, R, S V Gordon, M Marmiesse, P Brodin, C Buchrieser, K Eiglmeier, T Garnier, et al. 2002. “A New Evolutionary Scenario for the Mycobacterium Tuberculosis Complex.” Proceedings of the National Academy of Sciences 99 (6): 3684–89. https://doi.org/10.1073/pnas.052548299.

Camacho, Christiam, George Coulouris, Vahram Avagyan, Ning Ma, Jason Papadopoulos, Kevin Bealer, and Thomas L Madden. 2009. “BLAST+: Architecture and Applications.” BMC Bioinformatics 10 (December): 421. https://doi.org/10.1186/1471-2105-10-421.

Camus, Jean-Christophe, Melinda J Pryor, Claudine Me, and Stewart T Cole. 2019. “Re-Annotation of the Genome Sequence of Mycobacterium Tuberculosis H37Rv.” Vol. 24. www.sanger.ac.uk.

Cavet, Jennifer S., Alison I. Graham, Wenmao Meng, and Nigel J. Robinson. 2003. “A Cadmium-Lead-Sensing ArsR-SmtB Repressor with Novel Sensory Sites. Complementary Metal Discrimination by NmtR and CmtR in a Common Cytosol.” Journal of Biological Chemistry 278 (45): 44560–66. https://doi.org/10.1074/jbc.M307877200.

Chin, Chen Shan, David H. Alexander, Patrick Marks, Aaron A. Klammer, James Drake, Cheryl Heiner, Alicia Clum, et al. 2013. “Nonhybrid, Finished Microbial Genome Assemblies from Long-Read SMRT Sequencing Data.” Nature Methods 10 (6): 563–69. https://doi.org/10.1038/nmeth.2474.

Chiner-Oms, L. Sánchez-Busó, J. Corander, S. Gagneux, S. R. Harris, D. Young, F. González-Candelas, and I. Comas. 2019. “Genomic Determinants of Speciation and Spread of the Mycobacterium Tuberculosis Complex.” Science Advances 5 (6). https://doi.org/10.1126/sciadv.aaw3307.

Church, Deanna M., Valerie A. Schneider, Karyn Meltz Steinberg, Michael C. Schatz, Aaron R. Quinlan, Chen Shan Chin, Paul A. Kitts, et al. 2015. “Extending Reference Assembly Models.” Genome Biology 16 (1): 1–5. https://doi.org/10.1186/s13059-015-0587-3.

Cole, S. T., R. Brosch, J. Parkhill, T. Garnier, C. Churcher, D. Harris, S. V. Gordon, et al. 1998. “Deciphering the Biology of Mycobacterium Tuberculosis from the Complete Genome Sequence.” Nature 393 (6685): 537–44. https://doi.org/10.1038/31159.

Coll, Francesc, Ruth McNerney, José Afonso Guerra-Assunção, Judith R. Glynn, João Perdigão, Miguel Viveiros, Isabel Portugal, Arnab Pain, Nigel Martin, and Taane G. Clark. 2014. “A Robust SNP Barcode for Typing Mycobacterium Tuberculosis Complex Strains.” Nature Communications 5 (1): 4812. https://doi.org/10.1038/ncomms5812.

Comas, Iñaki, Mireia Coscolla, Tao Luo, Sonia Borrell, Kathryn E Holt, Midori Kato-Maeda, Julian Parkhill, et al. 2013. “Out-of-Africa Migration and Neolithic Coexpansion of Mycobacterium Tuberculosis with Modern Humans.” Nature Genetics 45 (10). https://doi.org/10.1038/ng.2744.

Coscolla, M, Daniela Brites, Fabrizio Menardo, Chloe Loiseau, Isaac Darko Otchere, Adwoa Asante-Poku, Prince Asare, et al. 2020. “Phylogenomics of Mycobacterium Africanum Reveals a New Lineage and a Complex Evolutionary History.” BioRxiv 17 (June): 19. https://doi.org/10.1101/2020.06.10.141788.

Coscolla, Mireia, and Sebastien Gagneux. 2014. “Consequences of Genomic Diversity in Mycobacterium Tuberculosis.” Seminars in Immunology 26 (6): 431–44. https://doi.org/10.1016/j.smim.2014.09.012.

Darko Otchere, Isaac, Mireia Coscollá, Leonor Sánchez-Busó, Adwoa Asante-Poku, Daniela Brites, Chloe Loiseau, Conor Meehan, et al. 2018. “Comparative Genomics of Mycobacterium Africanum Lineage 5 and Lineage 6 from Ghana Suggests Distinct Ecological Niches.” SCIENTIfIC RepoRtS | 8: 11269. https://doi.org/10.1038/s41598-018-29620-2.

Darling, Aaron C E, Bob Mau, Frederick R. Blattner, and Nicole T. Perna. 2004. “Mauve: Multiple Alignment of Conserved Genomic Sequence with Rearrangements.” Genome Research 14 (7): 1394–1403. https://doi.org/10.1101/gr.2289704.

DeJong, Bouke C, Philip C Hill, Alex Aiken, Timothy Awine, Martin Antonio, Ifedayo M Adetifa, Dolly J Jackson-Sillah, et al. 2008. “Progression to Active Tuberculosis, but Not Transmission, Varies by M. Tuberculosis Lineage in The Gambia.” J Infect Dis 198 (7): 1037–43. https://doi.org/10.1086/591504.

Dick, Thomas, Ujjini Manjunatha, Barbara Kappes, and Martin Gengenbacher. 2010. “Vitamin B6 Biosynthesis Is Essential for Survival and Virulence of Mycobacterium Tuberculosis.” Molecular Microbiology 78 (4): 980–88. https://doi.org/10.1111/j.1365-2958.2010.07381.x.

Firdessa, Rebuma, Stefan Berg, Elena Hailu, Esther Schelling, Balako Gumi, Girume Erenso, Endalamaw Gadisa, et al. 2013. “Mycobacterial Lineages Causing Pulmonary and Extrapulmonary Tuberculosis, Ethiopia.” Emerging Infectious Diseases 19 (3): 460–63. https://doi.org/10.3201/eid1903.120256.

Gagneux, Sebastien. 2018. “Ecology and Evolution of Mycobacterium Tuberculosis.” Nature Reviews Microbiology. https://doi.org/10.1038/nrmicro.2018.8.

Gagneux, Sebastien, Kathryn Deriemer, Tran Van, Midori Kato-Maeda, Bouke C De Jong, Sujatha Narayanan, Mark Nicol, et al. 2006. “Variable Host-Pathogen Compatibility in Mycobacterium Tuberculosis.” www.pnas.orgcgidoi10.1073pnas.0511240103.

Gao, Juehua, Jie Liao, and Guang-Yu Yang. 2009a. “Review Article CAAX-Box Protein, Prenylation Process and Carcinogenesis.” Am J Transl Res. Vol. 1. www.ajtr.org/AJTR905003.

Gao, Juehua, Jie Liao, and Guang Yu Yang. 2009b. “CAAX-Box Protein, Prenylation Process and Carcinogenesis.” American Journal of Translational Research.

Garimella, Kiran V, Zamin Iqbal, Michael A. Krause, Susana Campino, Mihir Kekre, Eleanor Drury, Dominic Kwiatkowski, Juliana M. Sa, Thomas E. Wellems, and Gil McVean. 2019. “Detection of Simple and Complex de Novo Mutations without, with, or with Multiple Reference Sequences.” BioRxiv, July, 698910. https://doi.org/10.1101/698910.

Gehre, Florian, Jacob Otu, Kathryn DeRiemer, Paola Florez de Sessions, Martin L. Hibberd, Wim Mulders, Tumani Corrah, Bouke C. de Jong, and Martin Antonio. 2013. “Deciphering the Growth Behaviour of Mycobacterium Africanum.” PLoS Neglected Tropical Diseases 7 (5). https://doi.org/10.1371/journal.pntd.0002220.

Ghodousi, Arash, Alamdar Hussain Rizvi, Aurangzaib Quadir Baloch, Abdul Ghafoor, Faisal Masood, Mehmood Qadir, Emanuele Borroni, Alberto Trovato, Sabira Tahseen, and Daniela Maria. 2019. “Acquisition of Cross-Resistance to Bedaquiline and Clofazimine Following Treatment for Tuberculosis in 2 Pakistan.” https://doi.org/10.1101/630012.

Goig, Galo A., Silvia Blanco, Alberto L. Garcia-Basteiro, and Iñaki Comas. 2020. “Contaminant DNA in Bacterial Sequencing Experiments Is a Major Source of False Genetic Variability.” BMC Biology 18 (1): 24. https://doi.org/10.1186/s12915-020-0748-z.

Goig, Galo A, Silvia Blanco, Alberto L. Garcia-Basteiro, and Iñaki Comas. 2019. “Contaminant DNA in Bacterial Sequencing Experiments Is a Major Source of False Genetic Variability.” BioRxiv.

Goig, Galo A, Silvia Blanco, Alberto L Garcia-Basteiro, and Iñaki Comas. 2018. “Pervasive Contaminations in Sequencing Experiments Are a Major Source of False Genetic Variability: A Mycobacterium Tuberculosis Meta-Analysis.” https://doi.org/10.1101/403824.

Gonzalo-Asensio, Jesús, Wladimir Malaga, Alexandre Pawlik, Catherine Astarie-Dequeker, Charlotte Passemar, Flavie Moreau, Françoise Laval, et al. 2014. “Evolutionary History of Tuberculosis Shaped by Conserved Mutations in the PhoPR Virulence Regulator.” Proceedings of the National Academy of Sciences of the United States of America 111 (31): 11491–96. https://doi.org/10.1073/pnas.1406693111.

Gordon, Stephen V., Roland Brosch, Alain Billault, Thierry Garnier, Karin Eiglmeier, and Stewart T. Cole. 1999. “Identification of Variable Regions in the Genomes of Tubercle Bacilli Using Bacterial Artificial Chromosome Arrays.” Molecular Microbiology 32 (3): 643–55. https://doi.org/10.1046/j.1365-2958.1999.01383.x.

Grinter, Rhys, Blair Ney, Rajini Brammananth, Christopher K. Barlow, Paul R.F. Cordero, David L. Gillett, Thierry Izoré, et al. 2020. “Cellular and Structural Basis of Synthesis of the Unique Intermediate Dehydro-F420-0 in Mycobacteria.” BioRxiv, February, 2020.02.27.968891. https://doi.org/10.1101/2020.02.27.968891.

Gu, J, H Li, M Li, C Vuong, M Otto, Y Wen, and Q Gao. 2005. “Bacterial Insertion Sequence IS256 as a Potential Molecular Marker to Discriminate Invasive Strains from Commensal Strains of Staphylococcus Epidermidis.” The Journal of Hospital Infection 61 (4): 342–48. https://doi.org/10.1016/j.jhin.2005.04.017.

Herrmann, Jean Louis, Robin Delahay, Alex Gallagher, Brian Robertson, and Douglas Young. 2000. “Analysis of Post-Translational Modification of Mycobacterial Proteins Using a Cassette Expression System.” FEBS Letters 473 (3): 358–62. https://doi.org/10.1016/S0014-5793(00)01553-2.

Ingen, Jakko van, Thomas A. Kohl, Katharina Kranzer, Barbara Hasse, Peter M. Keller, Anna Katarzyna Szafrańska, Doris Hillemann, et al. 2017. “Global Outbreak of Severe Mycobacterium Chimaera Disease after Cardiac Surgery: A Molecular Epidemiological Study.” The Lancet Infectious Diseases 17 (10): 1033–41. https://doi.org/10.1016/S1473-3099(17)30324-9.

Intemann, Christopher D., Thorsten Thye, Stefan Niemann, Edmund N. L. Browne, Margaret Amanua Chinbuah, Anthony Enimil, John Gyapong, et al. 2009. “Autophagy Gene Variant IRGM −261T Contributes to Protection from Tuberculosis Caused by Mycobacterium Tuberculosis but Not by M. Africanum Strains.” Edited by William Bishai. PLoS Pathogens 5 (9): e1000577. https://doi.org/10.1371/journal.ppat.1000577.

Iqbal, Zamin, Mario Caccamo, Isaac Turner, Paul Flicek, and Gil Mcvean. 2012. “De Novo Assembly and Genotyping of Variants Using Colored de Bruijn Graphs.” https://doi.org/10.1038/ng.1028.

Jandrasits, Christine, Stefan Kröger, Walter Haas, and Bernhard Y. Renard. 2019. “Computational Pan-Genome Mapping and Pairwise SNP-Distance Improve Detection of Mycobacterium Tuberculosis Transmission Clusters.” Edited by Sergei L. Kosakovsky Pond. PLOS Computational Biology 15 (12): e1007527. https://doi.org/10.1371/journal.pcbi.1007527.

Jong, Bouke C De, Martin Antonio, and Sebastien Gagneux. n.d. “Mycobacterium Africanum-Review of an Important Cause of Human Tuberculosis in West Africa.” Accessed May 21, 2019. https://doi.org/10.1371/journal.pntd.0000744.

Kalyaanamoorthy, Subha, Bui Quang Minh, Thomas K F Wong, Arndt von Haeseler, and Lars S Jermiin. 2017. “ModelFinder: Fast Model Selection for Accurate Phylogenetic Estimates.” Nature Methods 14 (6): 587–89. https://doi.org/10.1038/nmeth.4285.

Kapopoulou, Adamandia, Jocelyne M. Lew, and Stewart T. Cole. 2011. “The MycoBrowser Portal: A Comprehensive and Manually Annotated Resource for Mycobacterial Genomes.” Tuberculosis 91 (1): 8–13. https://doi.org/10.1016/j.tube.2010.09.006.

Kato-Maeda, Midori, Jeanne T Rhee, Thomas R Gingeras, Hugh Salamon, Jorg Drenkow, Nat Smittipat, and Peter M Small. 2001. “Comparing Genomes within the Species Mycobacterium Tuberculosis.” https://doi.org/10.1101/gr166401.

Kavanagh, K L, H Jçrnvall, B Persson, and U Oppermann. 2008. “The SDR Superfamily: Functional and Structural Diversity within a Family of Metabolic and Regulatory Enzymes” 171: 77. https://doi.org/10.1007/s00018-008-8588-y.

Kohl, Thomas Andreas, Christian Utpatel, Viola Schleusener, Maria Rosaria De Filippo, Patrick Beckert, Daniela Maria Cirillo, and Stefan Niemann. 2018. “MTBseq: A Comprehensive Pipeline for Whole Genome Sequence Analysis of Mycobacterium Tuberculosis Complex Isolates.” https://doi.org/10.7717/peerj.5895.

Leao, Sylvia, Anandi Martin, Gloria Isabel Mejia, Francoise Portaels, and Por-Taels F Leao SC, Martin A, Mejia GI, Palomino JC, Robledo J, Telles MAS. 2004. “Practical Handbook for the Phenotypic and Genotypic Identification of Mycobacteria.,” 77–126. https://doi.org/10.1007/BF01045823.

Lee, Robyn S., Jean François Proulx, Fiona McIntosh, Marcel A. Behr, and William P. Hanage. 2020. “Previously Undetected Super-Spreading of Mycobacterium Tuberculosis Revealed by Deep Sequencing.” ELife 9 (February). https://doi.org/10.7554/eLife.53245.

Lee, Robyn S, and Marcel A Behr. 2016. “Does Choice Matter? Reference-Based Alignment for Molecular Epidemiology of Tuberculosis.” Journal of Clinical Microbiology 54 (7): 1891–95. https://doi.org/10.1128/JCM.00364-16.

Lew, Jocelyne M., Adamandia Kapopoulou, Louis M. Jones, and Stewart T. Cole. 2011. “TubercuList - 10 Years After.” Tuberculosis 91 (1): 1–7. https://doi.org/10.1016/j.tube.2010.09.008.

Li, Ruiqiang, Yingrui Li, Hancheng Zheng, Ruibang Luo, Hongmei Zhu, Qibin Li, Wubin Qian, et al. 2010. “Building the Sequence Map of the Human Pan-Genome.” Nature Biotechnology 28 (1): 57–62. https://doi.org/10.1038/nbt.1596.

Magnus, K. 1966. “Epidemiological Basis of Tuberculosis Eradication 3. Risk of Pulmonary Tuberculosis after Human and Bovine Infection *.” Bull. Org. Mond. Sante. Vol. 35. https://www.ncbi.nlm.nih.gov/pmc/articles/PMC2476032/pdf/bullwho00607-0027.pdf.

Malone, Kerri M, Damien Farrell, Tod P Stuber, Olga T Schubert, Ruedi Aebersold, Suelee Robbe-Austerman, and Stephen V Gordon. 2017. “Updated Reference Genome Sequence and Annotation of Mycobacterium Bovis AF2122/97.” Genome Announcements 5 (14): e00157–17. https://doi.org/10.1128/genomeA.00157-17.

Marcus, Sarah A, Sarah W Sidiropoulos, Howard Steinberg, and Adel M Talaat. 2016. “CsoR Is Essential for Maintaining Copper Homeostasis in Mycobacterium Tuberculosis.” https://doi.org/10.1371/journal.pone.0151816.

Maretty, Lasse, Jacob Malte Jensen, Bent Petersen, Jonas andreas sibbesen, siyang Liu, Palle Villesen, Laurits skov, et al. 2017. “Sequencing and de Novo Assembly of 150 Genomes from Denmark as a Population Reference.” Nature Publishing Group 548. https://doi.org/10.1038/nature23264.

Marschall, Tobias, Manja Marz, Thomas Abeel, Louis Dijkstra, Bas E. Dutilh, Ali Ghaffaari, Paul Kersey, et al. 2018. “Computational Pan-Genomics: Status, Promises and Challenges.” Briefings in Bioinformatics 19 (1): 118–35. https://doi.org/10.1093/bib/bbw089.

Martiniano, Rui, Erik Garrison, Eppie R Jones, Andrea Manica, and Richard Durbin. 2019. “Removing Reference Bias in Ancient DNA Data Analysis by Mapping to a Sequence Variation Graph.” BioRxiv, September, 782755. https://doi.org/10.1101/782755.

McInerney, James O., Fiona J. Whelan, Maria Rosa Domingo-Sananes, Alan McNally, and Mary J. O’Connell. 2020. “Pangenomes and Selection: The Public Goods Hypothesis.” In The Pangenome, 151–67. Springer International Publishing. https://doi.org/10.1007/978-3-030-38281-0_7.

Medini, Duccio, Claudio Donati, Hervé Tettelin, Vega Masignani, and Rino Rappuoli. 2005. “The Microbial Pan-Genome.” Current Opinion in Genetics and Development. Elsevier Current Trends. https://doi.org/10.1016/j.gde.2005.09.006.

Meehan, Conor J., Pieter Moris, Thomas A. Kohl, Jūlija Pečerska, Suriya Akter, Matthias Merker, Christian Utpatel, et al. 2018. “The Relationship between Transmission Time and Clustering Methods in Mycobacterium Tuberculosis Epidemiology.” EBioMedicine 37 (November): 410–16. https://doi.org/10.1016/j.ebiom.2018.10.013.

Meehan, Conor J, Galo A Goig, Thomas A Kohl, Lennert Verboven, Anzaan Dippenaar, Matthew Ezewudo, Maha R Farhat, et al. 2019. “Whole Genome Sequencing of Mycobacterium Tuberculosis: Current Standards and Open Issues.” Nature Reviews Microbiology. https://doi.org/10.1038/s41579-019-0214-5.

Merker, Matthias, Thomas A Kohl, Ivan Barilar, Sönke Andres, Philip W Fowler, Erja Chryssanthou, Kristian Ängeby, et al. 2020. “Phylogenetically Informative Mutations in Genes Implicated in Antibiotic Resistance in Mycobacterium Tuberculosis Complex.” Genome Medicine. https://doi.org/10.1186/s13073-020-00726-5.

Merle, Corinne S.C., Charalambos Sismanidis, Oumou B. Sow, Martin Gninafon, John Horton, Olivier Lapujade, Mame B. Lo, et al. 2012. “A Pivotal Registration Phase III, Multicenter, Randomized Tuberculosis Controlled Trial: Design Issues and Lessons Learnt from the Gatifloxacin for TB (OFLOTUB) Project.” Trials 13 (May). https://doi.org/10.1186/1745-6215-13-61.

Mostowy, Serge, Debby Cousins, Jacqui Brinkman, Alicia Aranaz, and Marcel A Behr. 2002. “Genomic Deletions Suggest a Phylogeny for the Mycobacterium Tuberculosis Complex.” JID.

Murugesan, Saravanan, Stalin Mani, Indhumathy Kuppusamy, and Padma Krishnan. 2018. “Role of Insertion Sequence Element Is256 as a Virulence Marker and Its Association with Biofilm Formation among Methicillin-Resistant Staphylococcus Epidermidis from Hospital and Community Settings in Chennai, South India.” Indian Journal of Medical Microbiology 36 (1): 124. https://doi.org/10.4103/ijmm.IJMM_17_276.

Nebenzahl-Guimaraes, Hanna, Solomon A. Yimer, Carol Holm-Hansen, Jessica De Beer, Roland Brosch, and Dick Van Soolingen. 2016. “Genomic Characterization of Mycobacterium Tuberculosis Lineage 7 and a Proposed Name: ‘Aethiops Vetus.’” Microbial Genomics 2 (6). https://doi.org/10.1099/mgen.0.000063.

Ngabonziza, Jean Claude Semuto, Chloé Loiseau, Michael Marceau, Agathe Jouet, Fabrizio Menardo, Oren Tzfadia, Esdras Belamo Niyigena, et al. 2020. A Sister Lineage of the Mycobacterium Tuberculosis Complex Discovered in the African Great Lakes Region. BioRxiv. https://doi.org/10.1101/2020.01.20.912998.

Niemann, S., T. Kubica, F. C. Bange, O. Adjei, E. N. Browne, M. A. Chinbuah, R. Diel, et al. 2004. “The Species Mycobacterium Africanum in the Light of New Molecular Markers.” Journal of Clinical Microbiology 42 (9): 3958–62. https://doi.org/10.1128/JCM.42.9.3958-3962.2004.

Norman, Anders, Dorte Bek Folkvardsen, Søren Overballe-Petersen, and Troels Lillebaek. 2019. “Complete Genome Sequence of Mycobacterium Tuberculosis DKC2, the Predominant Danish Outbreak Strain.” https://doi.org/10.1128/MRA.

O’Toole, Ronan F., and Sanjay S. Gautam. 2017. “Limitations of the Mycobacterium Tuberculosis Reference Genome H37Rv in the Detection of Virulence-Related Loci.” Genomics 109 (5–6): 471–74. https://doi.org/10.1016/j.ygeno.2017.07.004.

Ofori-Anyinam, Boatema, Fatoumatta Kanuteh, Schadrac C. Agbla, Ifedayo Adetifa, Catherine Okoi, Gregory Dolganov, Gary Schoolnik, et al. 2016. “Impact of the Mycobaterium Africanum West Africa 2 Lineage on TB Diagnostics in West Africa: Decreased Sensitivity of Rapid Identification Tests in The Gambia.” PLoS Neglected Tropical Diseases 10: 1–12. https://doi.org/10.1371/journal.pntd.0004801.

Ofori-Anyinam, Boatema, Abi Janet Riley, Tijan Jobarteh, Ensa Gitteh, Binta Sarr, Tutty Isatou Faal-Jawara, Leen Rigouts, et al. 2020. “Comparative Genomics Shows Differences in the Electron Transport and Carbon Metabolic Pathways of Mycobacterium Africanum Relative to Mycobacterium Tuberculosis and Suggests an Adaptation to Low Oxygen Tension.” Tuberculosis 120 (January): 101899. https://doi.org/10.1016/j.tube.2020.101899.

Ondov, Brian D., Todd J. Treangen, Páll Melsted, Adam B. Mallonee, Nicholas H. Bergman, Sergey Koren, and Adam M. Phillippy. 2016. “Mash: Fast Genome and Metagenome Distance Estimation Using MinHash.” Genome Biology 17 (1): 132. https://doi.org/10.1186/s13059-016-0997-x.

Paten, Benedict, Adam M. Novak, Jordan M. Eizenga, and Erik Garrison. 2017. “Genome Graphs and the Evolution of Genome Inference.” Genome Research. Cold Spring Harbor Laboratory Press. https://doi.org/10.1101/gr.214155.116.

Periwal, Vinita, Ashok Patowary, Shamsudheen Karuthedath Vellarikkal, Anju Gupta, Meghna Singh, Ashish Mittal, Shamini Jeyapaul, et al. 2015. “Comparative Whole-Genome Analysis of Clinical Isolates Reveals Characteristic Architecture of Mycobacterium Tuberculosis Pangenome.” https://doi.org/10.1371/journal.pone.0122979.

Phelan, Jody, Paola Florez De Sessions, Leopold Tientcheu, Joao Perdigao, Diana Machado, Rumina Hasan, Zahra Hasan, et al. 2018. “Methylation in Mycobacterium Tuberculosis Is Lineage Specific with Associated Mutations Present Globally.” Scientific Reports 8 (1): 1–7. https://doi.org/10.1038/s41598-017-18188-y.

Rademacher, Corinna, and Bernd Masepohl. 2012. “Copper-Responsive Gene Regulation in Bacteria.” https://doi.org/10.1099/mic.0.058487-0.

Rakocevic, Goran, Vladimir Semenyuk, Wan Ping Lee, James Spencer, John Browning, Ivan J. Johnson, Vladan Arsenijevic, et al. 2019. “Fast and Accurate Genomic Analyses Using Genome Graphs.” Nature Genetics 51 (2): 354–62. https://doi.org/10.1038/s41588-018-0316-4.

Reiling, Norbert, Susanne Homolka, Kerstin Walter, Julius Brandenburg, Lisa Niwinski, Martin Ernst, Christian Herzmann, et al. 2013. “Clade-Specific Virulence Patterns of Mycobacterium Tuberculosis Complex Strains in Human Primary Macrophages and Aerogenically Infected Mice.” MBio 4 (4). https://doi.org/10.1128/mBio.00250-13.

Rengarajan, Jyothi, Barry R Bloom, and Eric J Rubin. 2005. “Genome-Wide Requirements for Mycobacterium Tuberculosis Adaptation and Survival in Macrophages.” PNAS June. Vol. 7. www.pnas.orgcgidoi10.1073pnas.0503272102.

Riojas, Marco A, Katya J Mcgough, Cristin J Rider-Riojas, Nalin Rastogi, and Manzour Hernando Hazbón. 2018. “Phylogenomic Analysis of the Species of the Mycobacterium Tuberculosis Complex Demonstrates That Mycobacterium Africanum, Mycobacterium Bovis, Mycobacterium Caprae, Mycobacterium Microti and Mycobacterium Pinnipedii Are Later Heterotypic Synonyms of Mycob.” https://doi.org/10.1099/ijsem.0.002507.

Rowland, Jennifer L, and Michael Niederweis. 2012. “Resistance Mechanisms of Mycobacterium Tuberculosis against Phagosomal Copper Overload.” https://doi.org/10.1016/j.tube.2011.12.006.

S.R. Pattyn, F. Portaels, L. Spanoghe, and J. Magos. 1970. “Further Studies on African Strains of Mycobacterium Tuberculosis. Comparison with M. Bovis and M. Microti.” Ann. Soc. belge Méd. Trop. https://www.ncbi.nlm.nih.gov/pubmed/4996738.

Samanovic, Marie I., Chen Ding, Dennis J. Thiele, and K. Heran Darwin. 2012. “Copper in Microbial Pathogenesis: Meddling with the Metal.” Cell Host & Microbe 11 (2): 106–15. https://doi.org/10.1016/j.chom.2012.01.009.

Sanoussi, C. N’Dira, Dissou Affolabi, Leen Rigouts, Séverin Anagonou, and Bouke de Jong. 2017. “Genotypic Characterization Directly Applied to Sputum Improves the Detection of Mycobacterium Africanum West African 1, under-Represented in Positive Cultures.” PLOS Neglected Tropical Diseases 11 (9): e0005900. https://doi.org/10.1371/journal.pntd.0005900.

Sanoussi, C. N’Dira, Bouke C De Jong, Mathieu Odoun, Karamatou Arekpa, Moulikatou Ali Ligali, Ousman Bodi, Simon Harris, et al. 2018. “Low Sensitivity of the MPT64 Identification Test to Detect Lineage 5 of the Mycobacterium Tuberculosis Complex.” https://doi.org/10.1099/jmm.0.000846.

Seemann, T. 2014. “Prokka: Rapid Prokaryotic Genome Annotation.” Bioinformatics 30 (14): 2068–69. https://doi.org/10.1093/bioinformatics/btu153.

Selengut, Jeremy D, and Daniel H Haft. 2010. “Unexpected Abundance of Coenzyme F 420-Dependent Enzymes in Mycobacterium Tuberculosis and Other Actinobacteria †.” JOURNAL OF BACTERIOLOGY 192 (21): 5788–98. https://doi.org/10.1128/JB.00425-10.

Sellés Vidal, Lara, Ciarán L. Kelly, Paweł M. Mordaka, and John T. Heap. 2018. “Review of NAD(P)H-Dependent Oxidoreductases: Properties, Engineering and Application.” Biochimica et Biophysica Acta (BBA) - Proteins and Proteomics 1866 (2): 327–47. https://doi.org/10.1016/j.bbapap.2017.11.005.

Sherman, Rachel M., Juliet Forman, Valentin Antonescu, Daniela Puiu, Michelle Daya, Nicholas Rafaels, Meher Preethi Boorgula, et al. 2019. “Assembly of a Pan-Genome from Deep Sequencing of 910 Humans of African Descent.” Nature Genetics. Nature Publishing Group. https://doi.org/10.1038/s41588-018-0273-y.

Szklarczyk, Damian, John H. Morris, Helen Cook, Michael Kuhn, Stefan Wyder, Milan Simonovic, Alberto Santos, et al. 2017. “The STRING Database in 2017: Quality-Controlled Protein-Protein Association Networks, Made Broadly Accessible.” Nucleic Acids Research 45 (D1): D362–68. https://doi.org/10.1093/nar/gkw937.

Tatusov, Roman L, Michael Y Galperin, Darren A Natale, and Eugene V Koonin. 2000. “The COG Database: A Tool for Genome-Scale Analysis of Protein Functions and Evolution.” Nucleic Acids Research. Vol. 28. http://www.ncbi.nlm.nih.gov/COG.

Tettelin, Hervé, Vega Masignani, Michael J. Cieslewicz, Claudio Donati, Duccio Medini, Naomi L. Ward, Samuel V. Angiuoli, et al. 2005. “Genome Analysis of Multiple Pathogenic Isolates of Streptococcus Agalactiae: Implications for the Microbial ‘Pan-Genome.’” Proceedings of the National Academy of Sciences of the United States of America 102 (39): 13950–55. https://doi.org/10.1073/pnas.0506758102.

Thye, Thorsten, Stefan Niemann, Kerstin Walter, Susanne Homolka, Christopher D. Intemann, Margaret Amanua Chinbuah, Anthony Enimil, et al. 2011. “Variant G57E of Mannose Binding Lectin Associated with Protection against Tuberculosis Caused by Mycobacterium Africanum but Not by M. Tuberculosis.” PLoS ONE 6 (6). https://doi.org/10.1371/journal.pone.0020908.

Tsolaki, Anthony G., Sebastien Gagneux, Alexander S. Pym, Yves Olivier L. Goguet De La Salmoniere, Barry N. Kreiswirth, Dick Van Soolingen, and Peter M. Small. 2005. “Genomic Deletions Classify the Beijing/W Strains as a Distinct Genetic Lineage of Mycobacterium Tuberculosis.” Journal of Clinical Microbiology 43 (7): 3185–91. https://doi.org/10.1128/JCM.43.7.3185-3191.2005.

Walker, Timothy M, Camilla LC Ip, Ruth H Harrell, Jason T Evans, Georgia Kapatai, Martin J Dedicoat, David W Eyre, et al. 2013. “Whole-Genome Sequencing to Delineate Mycobacterium Tuberculosis Outbreaks: A Retrospective Observational Study.” The Lancet Infectious Diseases 13 (2): 137–46. https://doi.org/10.1016/S1473-3099(12)70277-3.

Ward, Sarah K, Elizabeth A Hoye, and Adel M Talaat. 2008. “The Global Responses of Mycobacterium Tuberculosis to Physiological Levels of Copper †.” JOURNAL OF BACTERIOLOGY 190 (8): 2939–46. https://doi.org/10.1128/JB.01847-07.

WHO. 2016. “THE SHORTER MDR-TB REGIMEN FEATURES OF THE SHORTER MDR-TB REGIMEN REGIMEN COMPOSITION 4-6 Km-Mfx-Pto-Cfz-Z-H High-Dose-E / 5 Mfx-Cfz-Z-E.” www.who.int/tb.

WHO. 2019. WHO Consolidated Guidelines on Drug-Resistant Tuberculosis Treatment. http://apps.who.int/bookorders.

Zhang, Shuo, Jiazhen Chen, Peng Cui, Wanliang Shi, Wenhong Zhang, and Ying Zhang. 2015. “Identification of Novel Mutations Associated with Clofazimine Resistance in Mycobacterium Tuberculosis.” https://doi.org/10.1093/jac/dkv150.

