## Supplemental for "*Mycobacterium tuberculosis* complex lineage 5 exhibits high levels of within-lineage genomic diversity and differing gene content compared to the type strain H37Rv"

**Table S1. Gene difference within the three L5 completed PacBio genomes from Benin, Nigeria or The Gambia, and between these three genomes and *M. tuberculosis* H37Rv**

| **Genes** | **Present in PcbL5Ben** | **Present in PcbL5Gam** | **Present in PcbL5Nig** | **Present in H37Rv** |
|---|---|---|---|---|
| **Unshared (present in that genome only)** | PcbL5Ben_01893,<br>PcbL5Ben_01894,<br>PcbL5Ben_01895,<br>PcbL5Ben_02043,<br>PcbL5Ben_03043 | PcbL5Gam_02028,<br>PcbL5Gam_03020 | PcbL5Nig_02001 | **H37Rv_02002,<br>H37Rv_02085,<br>H37Rv_02086,<br>H37Rv_02087,<br>H37Rv_02107,<br>H37Rv_02108,<br>H37Rv_02109,<br>H37Rv_02188,<br>H37Rv_02189** |
| **Absent in PcbL5Ben** | | PcbL5Gam_unshared | PcbL5Nig_unshared | H37Rv_unshared |
| **Suspected pseudogenes in PcbL5Ben** | | PcbL5Gam_00036,<br>PcbL5Gam_00955,<br>PcbL5Gam_02045,<br>PcbL5Gam_03308,<br>PcbL5Gam_03376 | PcbL5Nig_00011,<br>PcbL5Nig_00036,<br>PcbL5Nig_00953,<br>PcbL5Nig_01794,<br>PcbL5Nig_01853,<br>PcbL5Nig_01896,<br>PcbL5Nig_02018,<br>PcbL5Nig_02119,<br>PcbL5Nig_02291,<br>PcbL5Nig_02467 | H37Rv_00012,<br>H37Rv_00036,<br>H37Rv_00528,<br>H37Rv_01371,<br>H37Rv_01801,<br>H37Rv_02020,<br>H37Rv_02463,<br>H37Rv_02556,<br>H37Rv_02951,<br>H37Rv_03349,<br>H37Rv_03797 |
| **Absent in PcbL5Gam** | PcbL5Ben_01443,<br>PcbL5Ben_01444,<br>PcbL5Ben_unshared | PcbL5Gam_unshared | PcbL5Nig_unshared | H37Rv_01413,<br>H37Rv_01414,<br>H37Rv_unshared |
| **Suspected pseudogenes in PcbL5Gam** | PcbL5Ben_01064,<br>PcbL5Ben_02378,<br>PcbL5Ben_02486,<br>PcbL5Ben_03113 | | PcbL5Nig_00011,<br>PcbL5Nig_01794,<br>PcbL5Nig_01853,<br>PcbL5Nig_01896,<br>PcbL5Nig_02119,<br>PcbL5Nig_02291,<br>PcbL5Nig_02467 | H37Rv_00012,<br>H37Rv_00528,<br>H37Rv_01371,<br>H37Rv_01801,<br>H37Rv_02329,<br>H37Rv_02463,<br>H37Rv_02556,<br>H37Rv_02951,<br>H37Rv_03797, |
| **Absent* in PcbL5Nig** | PcbL5Ben_01617,<br>PcbL5Ben_01618,<br>PcbL5Ben_01619,<br>PcbL5Ben_01620,<br>PcbL5Ben_01621, | PcbL5Gam_01604,<br>PcbL5Gam_01605,<br>PcbL5Gam_01606,<br>PcbL5Gam_01607,<br>PcbL5Gam_01608,<br>pcbL5Gam_01609, | | H37Rv_01581,<br>H37Rv_01582,<br>H37Rv_01583,<br>H37Rv_01584,<br>H37Rv_01585,<br>H37Rv_01586, |

| | | | | |
|---|---|---|---|---|
| | PcbL5Ben_01622,<br>PcbL5Ben_01623,<br>PcbL5Ben_01624,<br>PcbL5Ben_01625,<br>PcbL5Ben_01626,<br>PcbL5Ben_01627,<br>PcbL5Ben_01628,<br>PcbL5Ben_01629,<br>PcbL5Ben_01630,<br>PcbL5Ben_01631,<br>PcbL5Ben_01632,<br>PcbL5Ben_01633,<br>PcbL5Ben_01634,<br><br>PcbL5Ben_01636,<br>PcbL5Ben_01637,<br>PcbL5Ben_01638,<br>PcbL5Ben_01639,<br>PcbL5Ben_01640,<br>PcbL5Ben_01641,<br>PcbL5Ben_01642,<br>PcbL5Ben_01643,<br>PcbL5Ben_01644,<br>PcbL5Ben_01645,<br>PcbL5Ben_01646,<br>PcbL5Ben_01647,<br><br>PcbL5Ben_01891,<br>PcbL5Ben_01892,<br>PcbL5Ben_02248,<br><br>PcbL5Ben_unshared | PcbL5Gam_01610,<br>PcbL5Gam_01611,<br>PcbL5Gam_01612,<br>pcbL5Gam_01613,<br>PcbL5Gam_01614,<br>pcbL5Gam_01615,<br>PcbL5Gam_01616,<br>PcbL5Gam_01617,<br>PcbL5Gam_01618,<br>pcbL5Gam_01619,<br>PcbL5Gam_01620,<br>PcbL5Gam_01621,<br><br>PcbL5Gam_01623,<br>PcbL5Gam_01624,<br>PcbL5Gam_01625,<br>pcbL5Gam_01626,<br>PcbL5Gam_01627,<br>PcbL5Gam_01628,<br>PcbL5Gam_01629,<br>PcbL5Gam_01630,<br>PcbL5Gam_01631,<br>PcbL5Gam_01632,<br>pcbL5Gam_01633,<br>PcbL5Gam_01634,<br><br>PcbL5Gam_01881,<br>PcbL5Gam_01882,<br>PcbL5Gam_02228,<br>PcbL5Gam_unshared | | H37Rv_01587,<br>H37Rv_01588,<br>H37Rv_01589,<br>H37Rv_01590,<br>H37Rv_01591,<br>H37Rv_01592,<br>H37Rv_01593,<br>H37Rv_01594,<br>H37Rv_01595,<br>H37Rv_01596,<br>H37Rv_01597,<br>H37Rv_01598,<br><br>H37Rv_01600,<br>H37Rv_01601,<br>H37Rv_01602,<br>H37Rv_01603,<br>H37Rv_01604,<br>H37Rv_01605,<br>H37Rv_01606,<br>H37Rv_01607,<br>H37Rv_01608,<br>H37Rv_01609,<br>H37Rv_01610,<br><br>H37Rv_01853,<br>H37Rv_01854,<br>H37Rv_02202<br>H37Rv_unshared |
| **Suspected pseudogenes in PcbL5Nig** | PcbL5Ben_01064,<br>PcbL5Ben_01718,<br>PcbL5Ben_02486 | PcbL5Gam_01705,<br>PcbL5Gam_03308,<br>PcbL5Gam_03376 | | H37Rv_00528,<br>H37Rv_01371,<br>H37Rv_01679,<br>H37Rv_02463,<br>H37Rv_02556,<br>H37Rv_02951,<br>H37Rv_03349,<br>H37Rv_03797 |
| **Absent in H37Rv** | **PcbL5Ben_01364,<br>PcbL5Ben_01365,<br>PcbL5Ben_02129,<br>PcbL5Ben_02130,<br>PcbL5Ben_02181,** | **PcbL5Gam_01359,<br>PcbL5Gam_01360,<br>PcbL5Gam_02112,<br>PcbL5Gam_002113,<br>PcbL5Gam_02163,** | **PcbL5Nig_01357,<br>PcbL5Nig_01358,<br>PcbL5Nig_02085,<br>PcbL5Nig_02086,<br>PcbL5Nig_02137,** | |

| | | | |
|---|---|---|---|
| | **PcbL5Ben_02182,**<br>**PcbL5Ben_03555,**<br>**PcbL5Ben_03556,**<br>**PcbL5Ben_03557,**<br>**PcbL5Ben_03669**<br>PcbL5Ben_unshared | **PcbL5Gam_02164,**<br>**PcbL5Gam_03527,**<br>**PcbL5Gam_03528,**<br>**PcbL5Gam_03529,**<br>**PcbL5Gam_03643**,<br>PcbL5Gam_unshared | **PcbL5Nig_02138,**<br>**PcbL5Nig_03500,**<br>**PcbL5Nig_03503,**<br>**PcbL5Nig_03504,**<br>**PcbL5Nig_03616,**<br>PcbL5Nig_unshared |
| **Suspected pseudogenes in H37Rv** | PcbL5Ben_00749,<br>PcbL5Ben_00828,<br>PcbL5Ben_01064,<br>PcbL5Ben_01108,<br>PcbL5Ben_02128,<br>PcbL5Ben_02486,<br>PcbL5Ben_03113,<br>PcbL5Ben_03141,<br>PcbL5Ben_03667 | PcbL5Gam_00750,<br>PcbL5Gam_00826,<br>PcbL5Gam_00955,<br>PcbL5Gam_01103,<br>PcbL5Gam_01879,<br>PcbL5Gam_02111,<br>PcbL5Gam_03117,<br>PcbL5Gam_03308,<br>PcbL5Gam_03641 | PcbL5Nig_00748,<br>PcbL5Nig_00824,<br>PcbL5Nig_00953,<br>PcbL5Nig_01100,<br>PcbL5Nig_01846,<br>PcbL5Nig_01853,<br>PcbL5Nig_01896,<br>PcbL5Nig_02084,<br>PcbL5Nig_02119,<br>PcbL5Nig_02291,<br>PcbL5Nig_02467,<br>PcbL5Nig_03064,<br>PcbL5Nig_03092,<br>PcbL5Nig_03614 |

*Genes highlighted in the same color are contiguous and form a block.

**Table S2. Verification of genes in the L5 completed PacBio genomes suspected to be pseudogenes in *M. tuberculosis* H37Rv genome.**

| Gene | Gene in PcbL5Ben | Gene in PcbL5Gam | Gene in PcbL5Nig | Length (aa) | Locus in H37Rv | Remarks | Conclusion |
|---|---|---|---|---|---|---|---|
| **1** | **PcbL5Ben_00749** | **PcbL5Gam_00750** | **PcbL5Nig_00748** | 101 | 798857-799150 | M1V, L96p, H97t, del NCD (aa 99-101) | **Pseudogene in H37Rv** |
| **2** | PcbL5Ben_00828 | PcbL5Gam_00826 | PcbL5Nig_00824 | 34 | 863155-863256 | No change | Not pseudogene in H37Rv |
| **3** | Not suspected pseudogene | PcbL5Gam_00955 | PcbL5Nig_00953 | 204 | 993212-992602 | Inverted, M1V | **Pseudogene in H37Rv** |
| **4** | PcbL5Ben_01064 | Not suspected pseudogene | Not suspected pseudogene | 34 | 1102546-1102647 | S23r | Not pseudogene in H37Rv |
| **5** | **PcbL5Ben_01108** | **PcbL5Gam_01103** | **PcbL5Nig_01100** | 70 | 1148218-1148397 | M1V | **Pseudogene in H37Rv** |
| **6** | **PcbL5Ben_01889** | **PcbL5Gam_01879** | **PcbL5Nig_01846** | 68 | 1981341-1981341 | M1V | **Pseudogene in H37Rv** |
| **7** | **PcbL5Ben_02128** | **PcbL5Gam_02111** | **PcbL5Nig_02084** | 92 | 2219418-2219318 | Inverted, alignment showed aa 59-92, no mutation | **Pseudogene in H37Rv** |
| **8** | Not suspected pseudogene | Not suspected pseudogene | PcbL5Nig_01853 is PcbNig_1896 | 41 | 2038813-2038706 | Inverted, alignment showed aa 59-92, no mutation | **Pseudogene in H37Rv** |

| | | | | | | | |
|---|---|---|---|---|---|---|---|
| **9** | PcbL5Ben_02486 | Not suspected pseudogene | Not suspected pseudogene | 41 | 2582440-2582318 | Inverted, M1V | **Pseudogene in H37Rv** |
| **10** | **PcbL5Ben_03141** | **PcbL5Gam_03117** | **PcbL5Nig_03092** | 111 | 3291532-3291350 | Inverted, del aa 1-aa60 (60 first aa) | **Pseudogene in H37Rv** |
| **11** | Not suspected pseudogene | Not suspected pseudogene | PcbL5Nig_02119 | 46 | 2256620-2256483 | Inverted, No change | **Pseudogene in H37Rv** |
| **12** | Not suspected pseudogene | Not suspected pseudogene | PcbL5Nig_02291 | 33 | 24345567-2434665 | No change | Not pseudogene in H37Rv |
| **13** | Not suspected pseudogene | Not suspected pseudogene | PcbL5Nig_02467 | 94 | 2614352-2614071 | Inverted | **Pseudogene in H37Rv** |
| **14** | PcbL5Ben_03113 | Not suspected pseudogene | PcbL5Nig_03060 | 204 | 3232652-3232867 | Del aa1-aa56, N74D, del aa129-aa204 | **Pseudogene in H37Rv** |
| **15** | Not suspected pseudogene | PcbL5Gam_03308 | Not suspected pseudogene | 58 | 3487871-3487698 | Inverted, M1V | **Pseudogene in H37Rv** |
| **16** | **PcbL5Ben_03667** | **PcbL5Gam_03641** | **PcbL5Nig_03614** | 99 | 3841840-3842109 | M1V, all aa mutated (insertion substitution) from aa26-aa81, del aa82-aa98 | **Pseudogene in H37Rv** |

32

**Table S3. Verification of genes present in H37Rv suspected to be pseudogenes in the L5 completed genomes**

| Gene in H37Rv | Length (aa) | Locus in H37Rv | Locus in PcbL5Ben | Locus in PcbL5Gam | Locus in PcbL5Nig | Remarks | Conclusion |
|---|---|---|---|---|---|---|---|
| H37Rv_02463 | 42 | 2604950-2605078 | 2631987-2632112 | 2614961-2615086 | 2603370-2603495 | No change | Not pseudogene |
| **H37Rv_02556** | 67 | 2718844-2719047 | 2743020-2742820 | 2729937-2729737 | 2716988-2716788 | Inverted, nSNP I66L | **Pseudogene in the 3 Pacbio L5** |
| **H37Rv_02951** | 55 | 3115741-3115908 | 3137926-3137762 | 3126133-3125969 | 3112866-3112702 | Inverted, nSNP M1V, L55F | **Pseudogene in the 3 Pacbio L5** |
| **H37Rv_03797** | 200 | 4053036-4053638 | 4076665-4077105 | 4064716-4065156 | 4055914-4056354 | del aa 1-53 | **Pseudogene in the 3 Pacbio L5** |

**Table S4. Genes absent in the Nigerian completed PacBio genome (PcbL5Nig) and some L5 isolates Illumina genomes (n=6), but present in the Benin and Gambian completed PacBio genomes (PcbL5Ben, PcbL5Gam), H37Rv and all other L5 isolates Illumina genomes.**

| ID (this study) | Rv number | Length (bp) | Functional group (Mycobrowser) | Gene name (Mycobrowser) | Product (Mycobrowser) | Function (Mycobrowser) |
|---|---|---|---|---|---|---|
| H37Rv_01581 | *Rv1493* | 2253 bp | Lipid metabolism | *mutB* | Probable methylmalonyl-CoA mutase large subunit MutB (MCM) | Involved in propionic acid fermentation |
| H37Rv_01582 | *Rv1494* | 303 bp | Virulence, detoxification, adaptation | *mazE4* | Possible antitoxin MazE4 | Possible mazE4, antitoxin, part of toxin-antitoxin (TA) operon with Rv1495. Non-essential gene for in vitro growth of H37Rv |
| H37Rv_01583 | *Rv1495* | 318 bp | Virulence, detoxification, adaptation | *mazF4* | Possible toxin MazF4 | Sequence-specific mRNA cleavage |
| H37Rv_01584 | *Rv1496* | 1005 bp | Cell wall and cell processes | *Rv1496* | Possible transport system kinase | Possibly involved in transport (possibly arginine) |
| H37Rv_01585 | *Rv1497* | 1290 bp | Intermediary metabolism and respiration | *lipL* | Probable esterase LipL | Function unknown, but supposed involvement in lipid metabolism |
| H37Rv_01586 | *Rv1498A* | 213 bp | Conserved hypotheticals | *Rv1498A* | Conserved protein | Function unknown |
| H37Rv_01587 | *Rv1498c* | 618 bp | Intermediary metabolism and respiration | *Rv1498c* | Probable methyltransferase | Causes methylation |
| H37Rv_01588 | *Rv1499* | 399 bp | Conserved hypotheticals | *Rv1499* | Hypothetical protein | Function unknown |

| | | | | | | |
|---|---|---|---|---|---|---|
| H37Rv_01589 | *Rv1500* | 1029 bp | Intermediary metabolism and respiration | *Rv1500* | Probable glycosyltransferase | Function unknown |
| H37Rv_01590 | *Rv1501* | 822 bp | Conserved hypotheticals | *Rv1501* | Conserved hypothetical protein | Function unknown |
| H37Rv_01591 | *Rv1502* | 900 bp | Unknown | *Rv1502* | Hypothetical protein | Function unknown |
| H37Rv_01592 | *Rv1505c* | 666 bp | Conserved hypotheticals | *Rv1505c* | Conserved hypothetical protein | Function unknown.<br>It has some similarity to hypothetical proteins and glycosylases |
| H37Rv_01593 | *Rv1506c* | 501 bp | Unknown | *Rv1506c* | Hypothetical protein | Function unknown. Non-essential gene for in vitro growth of H37Rv |
| H37Rv_01594 | *Rv1507A* | 504 bp | | | | |
| H37Rv_01595 | *Rv1507c* | 696 bp | Conserved hypotheticals | *Rv1507A* | Hypothetical protein | Function unknown |
| H37Rv_01596 | *Rv1508A* | 636 bp | Conserved hypotheticals | *Rv1508A* | Conserved hypotheticals | Function unknown.<br>Highly similar to central part of glycosyl transferases from various mycobacteria and eubacteria |
| H37Rv_01597 | *Rv1508c* | 1800 bp | Cell wall and cell processes | *Rv1508c* | Probable membrane protein | Function unknown. Predicted to be in the GT-C superfamily of glycosyltransferases |
| H37Rv_01598 | *Rv1509* | 882 bp | Unknown | *Rv1509* | Hypothetical protein | Function unknown |
| H37Rv_01599 | *Rv1510* (present in | 1299 bp | Cell wall and cell processes | *Rv1510* | Probable conserved membrane protein | Function unknown |

| | | | | | | |
|---|---|---|---|---|---|---|
| | PcbL5Nig but missing in the PcbL5Nig-like Illumina L5) | | | | | |
| H37Rv_01600 | *Rv1511* | 1023 bp | Intermediary metabolism and respiration | *gmdA* | GDP-D-mannose dehydratase GmdA (GDP-mannose 4,6 dehydratase) | Function unknown, probably involved in nucleotide-sugar metabolism |
| H37Rv_01601 | *Rv1512* | 969 bp | Intermediary metabolism and respiration | *epiA* | Probable nucleotide-sugar epimerase EpiA | Function unknown, probably involved in nucleotide-sugar metabolism |
| H37Rv_01602 | *Rv1513* | 732 bp | Conserved hypotheticals | *Rv1513* | Conserved protein | Function unknown, similar to hypothetical proteins from several organisms |
| H37Rv_01603 | *Rv1514c* | 789 bp | Conserved hypotheticals | *Rv1514c* | Conserved hypothetical protein | Function unknown; similar to other hypothetical protein and to putative colanic acid biosynthesis glycosyl transferase |
| H37Rv_01604 | *Rv1515c* | 897 bp | Conserved hypotheticals | *Rv1515c* | Conserved hypothetical protein | Function unknown |
| H37Rv_01605 | *Rv1516c* | 1011 bp | Intermediary metabolism and respiration | *Rv1516c* | Probable sugar transferase | Function unknown, involved in cellular metabolism. Non essential for in vitro growth. |
| H37Rv_01606 | *Rv1517* | 765 bp | Cell wall and cell processes | *Rv1517* | Conserved hypothetical membrane protein | Function unknown |

| | | | | | | |
|---|---|---|---|---|---|---|
| H37Rv_01607 | *Rv1518* | 960 bp | Conserved hypotheticals | *Rv1518* | Conserved hypothetical protein | possibly glycosyl transferase involved in exopolysaccharide synthesis, similar to several hypothetical proteins and glycosyl transferases from diverse organisms |
| H37Rv_01608 | *Rv1519* | 270 bp | Conserved hypotheticals | *Rv1519* | Conserved hypothetical protein | Function unknown |
| H37Rv_01609 | *Rv1520* | 1041 bp | Intermediary metabolism and respiration | *Rv1520* | Probable sugar transferase | Function unknown; thought to be involved in cellular metabolism |
| H37Rv_01610 | *Rv1521* | 1752 bp | Lipid metabolism | *fadD25* | Probable fatty-acid-AMP ligase FadD25 (fatty-acid-AMP synthetase) (fatty-acid-AMP synthase) | Function unknown; involved in lipid degradation |
| | *Rv1522c* (present in PcbL5Nig but missing in the PcbL5Nig-like Illumina L5) | 3441 bp | Cell wall and cell processes | *mmpL12* | Probable conserved transmembrane transport protein MmpL12 | Function unknown. Thought to be involved in fatty acid transport. |

**Table S5. Presence /absence in 205 L5 isolates Illumina genomes of the 10 genes shared by the three L5 completed genomes but absent in the *M. tuberculosis* H37Rv genome.**

| Genes present in L5 completed PacBio genomes but absent in H37Rv | Length (bp) | Present in L5 isolates Illumina genome n=205 (%) | SNP in L5-specific genes | | | Gene function | | | Present in complete genome of |
|---|---|---|---|---|---|---|---|---|---|
| | | | PcbL5Ben as ref n=205 (%) | PcbL5Gam as ref n=205 (%) | PcbL5Nig as ref n=205 (%) | Functional group (Mycobrowser) | genes | Summary of role based on the literature | |
| **PcbL5Ben_1364** | 546 bp | 191 (93.2) | | | | ABC transporter ATP-binding protein/permease (Prokka) | Transmembrane ATP-binding protein ABC transporter | Conserved domains: FHA and CcmA (NCBI)<br>FHA (forkhead associated domain, COG1716 (T)) binds pSer, pThr, pTyr (Signal transduction mechanisms)<br>CcmA (COG1131 (V)), ABC-type multidrug transport system, ATPase | L5, bovis |

| | | | | | | | | | |
|---|---|---|---|---|---|---|---|---|---|
| | | | | | | | | component (Defense mechanisms (NCBI)) | |
| **PcbL5Ben_1365** | 1902 bp | 191 (93.2) | | | | Same as PcbL5Ben_1364 | Same as PcbL5Ben_1364 | Same as PcbL5Ben_1364 | L5, bovis |
| **PcbL5Ben_2129#** | 648 bp | **205 (100)** | 4 (2) | 4 (2) | 205* (100) | **PE/PPE** | PPE, possibly PE35 (77% id with PE35 from *M. braziliense*) | PE35 implicated in virulence (STRING database) and PE family is suggested to be related to antigenic variation of MTBC (NCBI) | **Only L5** |
| **PcbL5Ben_2130#** | 600 bp | **205 (100)** | 7 (3.4) | 7 (3.4) | 7 (3.4) | Not in Mycobrowser | **Hypothetical protein possibly CAAX conserved domain** (NCBI) | Involved in post-translation modification by attaching to the isoprenoid proteins in the process called prenylation (NCBI). Most of CAAX box proteins do not have a transmembrane domain, thus, the prenylation process is crucial for the function of many signal transduction proteins (Gao, Liao, and Yang 2009a) | **Only L5** |
| **PcbL5Ben_2181** | 237 bp | **205 (100)** | 3 (1.5) | 3 (1.5) | 3 (1.5) | Not in Mycobrowser | **Hypothetical protein, possibly IS256 transposase** (90% id, NCBI) | IS256 transposase is implicated in resistance to antibiotics and virulence (recombination for adaptation, invasion)(Murugesan et al. 2018; Gu et al. 2005) | L5, L6, bovis |
| **PcbL5Ben_2182** | 897 bp | **205 (100)** | 6 (2.9) | 6 (2.9) | 6 (2.9) | **Unknown function (Mb2048c**) | *Mb2048c* | Function unknown (expressed during exponential growth in Sauton's minimal media (NCBI)). Gene absent in H37Rv (Mycobrowser) | L5, L6, bovis |

| | | | | | | |
|---|---|---|---|---|---|---|
| **PcbL5Ben_3555** | 1137 bp | 167 (81.5) | Intermediary metabolism and respiration | moaA3 (NCBI) | *moaA3* (molybdenum cofactor biosynthesis protein subunit MoaA , Cyclic pyranopterin monophosphate synthase (Prokka)) is contained in an IS6110 sequence deleted in H37Rv (Mycobrowser) | L5, L6, bovis |
| **PcbL5Ben_3556** | 1146 bp | 165 (80.5) | Not in Mycobrowser | Hypothetical protein, possibly DnrI superfamily (NCBI) | *DnrI*: DNA binding transcriptional activator of the SARP family (signal transduction mechanisms, COG3629 (NCBI)) | L5, L6, bovis |
| **PcbL5Ben_3557** | 438 bp | 165 (80.5) | Peptide synthase (Prokka) | Hypothetical protein, possibly IS6110 transposase (99-100% id NCBI) | | L5, L6, bovis |
| **PcbL5Ben_3669** | 396 bp | 201 (98) | PE/PPE | Hypothetical protein | | L5, L6 |

Ref: reference ; bp: base pair

*In total 210 in the 205 L5 Illumina genomes: more than one SNP in the gene for some strains

### Present in L5 only, not present in L6 nor *M. bovis.* Those two genes (PcbL5Ben_2129 and 2130) are adjacent.

PcbL5Ben_1364 and PcbL5Ben_1365 are adjacent genes

PcbL5Ben_2181 and PcbL5Ben_2182 are adjacent genes

PcbL5Ben_3555, PcbL5Ben_3556 and PcbL5Ben_3557 are adjacent genes

**Table S6. Presence/absence in 205 Illumina L5 genomes of the nine genes present in the H37Rv genome but absent in the three completed L5 genomes.**

| Genes present in H37Rv but absent in L5 completed PacBio genomes | Length (bp) | Present in L5 isolates Illumina genomes<br>n=205 (%) | Absent in L5 isolates Illumina genomes<br>n=205 (%) | Gene function | | | Absent in complete genome of | Present in complete genome of |
|---|---|---|---|---|---|---|---|---|
| | | | | **Functional group (Mycobrowser)** | **Genes** | **Summary of role based on the literature** | | |
| **Part of Rv1899c (H37Rv_2002)** | 105 bp | 204 (99.5) | 1 (0.5) | Hypothetical protein (35 aa) part of *Rv1899c* (343 aa, Cell wall and cell processes)(Tuberculist) | Hypothetical protein, part of Rv1899c (LppD: 4-hydroxybuturate dehydrogenase 173 aa) and also LpqI beta-hexoaminidase precursor (Herrmann et al. 2000) | Possibly bacterial membrane lipoprotein (LppD and LpqI) | L5, L6, bovis | **H37Rv only** |
| **Rv1977 (H37Rv_2085)** | 1047 bp | **0** | **205 (100)** | **Conserved hypotheticals** | M48 peptidase family protein including as homologs:<br>CAAX prenyl protease (*Htpx* (COG0501 O, heat shock stress response protein)<br>M48-Ste24p-like | Bacterial survival (bacterial self-degradation to eliminate abnormal membrane proteins, htpx upregulated at high temperature, and also after 96 hours of starvation (NCBI)) | **L5 only** | L6, bovis, H37Rv |
| **Rv1978 (H37Rv_2086)** | 849 bp | 3 (1.5) | 202 (98.5) | Conserved hypotheticals, | M48-2C-Ste24p (100% id), class I SAM-dependent methyltransferase (99.65% id), type 11 methyltransferase (97.1% id) | Bacterial survival in macrophages, and non-essential for in vitro growth of H37Rv (Rengarajan, Bloom, and Rubin 2005)(Mycobrowser, Tuberculist) | L5 only | L6, bovis, H37Rv |

| | | | | | | | | |
|---|---|---|---|---|---|---|---|---|
| **Rv1979c (H37Rv_2087)** | 1446 bp | **0** | **205 (100)** | **Cell wall and cell processes** | APC family permease | -Involved in the transport of **clofazimine** (and **bedaquiline**), associated with clofazimine and bedaquiline resistance (Zhang et al. 2015; Ghodousi et al. 2019)-Disruption of the gene provides an in vitro growth advantage to H37Rv | **L5 only** | L6, bovis, H37Rv |
| **Rv1993c (H37Rv_2107)** | 273 bp | **0** | **205 (100)** | **Conserved hypotheticals** | *Rv0968 =DUF1490* similar to *Rv1993c* (NCBI) | Similar to *Rv0968* which is included in the operon *cosR-Rv0968-ctpV* where *cosR* and *ctpV* are virulence associated (Rademacher and Masepohl 2012) | **L5 only** | L6, bovis, H37Rv |
| **Rv1994c (H37Rv_2108)** | 357 bp | 1 (0.5) | 204 (99.5) | Regulatory proteins | HTH transcriptional regulator *cmtR* | Operon *cmtR-Rv1993c-ctpG* is similar to *csoR-Rv0968-ctpV*.<br>*cmtR* (*Rv1994c*) has the same role as *csoR*.<br>These operons all function in the regulation and transport (efflux) of toxic metal especially copper which is toxic in excess (Samanovic et al. 2012).<br>Disruption may hamper in vitro growth, and survival during chronic phase of infection as *csoR* absence (Marcus et al. 2016; Rowland and Niederweis 2012; Ward, Hoye, and Talaat 2008). | L5 only | L6, bovis, H37Rv |
| **Rv1995 (H37Rv_2109)** | 729 bp | **0** | **205 (100)** | **Conserved hypothetical** | **Hemerythrin-domain containing protein (NCBI)** | - Involved in oxygen transport (NCBI) | **L5 only** | L6, bovis, H37Rv |

| | | | | | | | | |
|---|---|---|---|---|---|---|---|---|
| **Rv2073c (H37Rv_2188)** | 750 bp | **0** | **205 (100)** | **Intermediary metabolism and respiration** | **Probable short chain dehydrogenase** | SDR family NAD(P)-dependent oxidoreductase (NCBI): catalyzes a wide range of reactions and substrate(Kavanagh et al. 2008; Sellés Vidal et al. 2018) | L5, L6, bovis | **H37Rv only** |
| **Rv2074 (H37Rv_2189)** | 408 bp | 1 (0.5) | 204 (99.5) | **Intermediary metabolism and respiration (Pyridoxamine, Vit B6)** | Pyridoxamine-5-phosphate oxidase (Mycobrowser, Tuberculist, NCBI)<br>Its cofactor (Selengut and Haft 2010) F420 dependent biliverdin reductase (99.2-100% id, NCBI)(Ahmed et al. 2016) | - Biosynthesis of (pyridoxal phosphate and) pyridoxine (vitamin B6, Mycobrowser) which is essential for survival and virulence of M. tuberculosis (Dick et al. 2010)<br>- Its cofactor implicated in Immuno-evasive mechanism to allow bacterial persistence (Ahmed et al. 2016; Selengut and Haft 2010) | L5, L6, bovis | H37Rv only |

**Table S7. Country of origin of 205 Illumina sequenced genomes**

| **L5 strains** | **Sublineage** |
|---|---|
| PcbL5Benin | L5.3 (L5.3.1) |
| PcbL5Gambia | L5.1.5 |
| PcbL5Nigeria | L5.3 (L5.3.2) |
| 10451-01 | L5.1.1 |
| 10463-02 | L5.1.1 |
| 10480-01 | L5.1.1 |
| 10519-01 | L5.1.1 |
| 10617-12 | L5.1.1 |
| 10695-13 | unknown (new sublineage) |
| 10709-13 | L5.1.1 |
| 11821-03 | unknown (new sublineage) |
| BSSE-QGF-22150 | L5.1.3 |
| BSSE-QGF-22151 | L5.1.1 |
| BSSE-QGF-22153 | L5.1.1 |
| BSSE-QGF-22154 | L5.1.4 |
| BSSE-QGF-22155 | L5.1.1 |
| BSSE-QGF-27970 | L5.1.1 |
| BSSE-QGF-28835 | L5.3 (L5.3.2) |
| BSSE-QGF-54153 | unknown (new sublineage) |
| BSSE-QGF-78571 | L5.1.4 |
| DRC-072384 | L5.2 |
| ERR019875 | L5.1.2 |
| ERR1023216 | unknown (new sublineage) |
| ERR1023217 | L5.1.3 |
| ERR1023218 | unknown (new sublineage) |
| ERR1023221 | L5.3 (L5.3.2) |
| ERR1023223 | L5.1.5 |
| ERR1023224 | L5.1.2 |
| ERR1082117 | L5.1.1 |
| ERR1082122 | L5.1.2 |
| ERR1082125 | L5.1.1 |
| ERR1082126 | L5.1.1 |
| ERR1082129 | L5.1.2 |
| ERR1082135 | L5.1.1 |
| ERR1082137 | L5.1.3 |
| ERR1203054 | L5.1.1 |
| ERR1203057 | L5.1.1 |
| ERR1203058 | L5.1.2 |
| ERR1203059 | L5.1.1 |
| ERR1203065 | L5.1.1 |
| ERR1203066 | L5.1.2 |
| 12026-12 | L5.1.1 |

| | |
|---|---|
| 1447-02 | unknown (new sublineage) |
| 4802-03 | L5.1.4 |
| 5438-02 | L5.1.1 |
| 7491-14 | L5.1.1 |
| BSSE-QGF-22149 | L5.1.2 |
| BSSE-QGF-54138 | L5.1.4 |
| ERR1082120 | L5.1.1 |
| ERR1203068 | L5.1.2 |
| ERR2383621 | L5.1.1 |
| ERR439939 | L5.1.4 |
| ERR439964 | L5.1.1 |
| ERR502505 | L5.1.1 |
| ERR702407 | L5.1.3 |
| ERR751308 | L5.1.1 |
| ERR751348 | L5.1.4 |
| ERR1203069 | L5.1.4 |
| ERR1203074 | L5.1.1 |
| ERR1215463 | unknown (new sublineage) |
| ERR1215473 | L5.1.5 |
| ERR1215476 | L5.1.4 |
| ERR1215477 | L5.1.1 |
| ERR1215478 | L5.1.2 |
| ERR1334053 | L5.1.1 |
| ERR234679 | L5.1.3 |
| ERR234680 | unknown (new sublineage) |
| ERR2383619 | L5.1.1 |
| ERR2383620 | L5.2 |
| ERR2383622 | L5.3 (L5.3.1) |
| ERR2383623 | L5.2 |
| ERR2383624 | L5.2 |
| ERR2383625 | L5.1.1 |
| ERR2383626 | L5.2 |
| ERR2704808 | L5.1.1 |
| ERR2704809 | L5.1.1 |
| ERR2704810 | L5.2 |
| ERR2704812 | L5.2 |
| ERR439931 | L5.1.3 |
| ERR439936 | L5.1.5 |
| ERR439937 | L5.1.2 |
| ERR439940 | L5.1.5 |
| ERR439941 | L5.1.4 |
| ERR439944 | unknown (new sublineage) |
| ERR439947 | L5.1.5 |
| ERR439949 | L5.1.4 |
| ERR439951 | L5.3 (L5.3.2) |

| | |
|---|---|
| ERR439952 | L5.3 (L5.3.1) |
| ERR439953 | L5.1.5 |
| ERR439955 | L5.1.4 |
| ERR439959 | L5.3 (L5.3.2) |
| ERR439960 | unknown (new sublineage) |
| ERR439962 | unknown (new sublineage) |
| ERR439967 | L5.1.1 |
| ERR439980 | L5.1.1 |
| ERR439982 | unknown (new sublineage) |
| ERR439983 | L5.3 (L5.3.1) |
| ERR439984 | unknown (new sublineage) |
| ERR439985 | unknown (new sublineage) |
| ERR460916 | L5.3 (L5.3.1) |
| ERR502471 | L5.1.1 |
| ERR502475 | L5.1.1 |
| ERR502487 | unknown (new sublineage) |
| ERR502500 | L5.1.1 |
| ERR502501 | L5.3 (L5.3.2) |
| ERR502509 | L5.1.5 |
| ERR502512 | L5.3 (L5.3.2) |
| ERR502513 | L5.1.4 |
| ERR502515 | L5.1.1 |
| ERR502536 | L5.1.4 |
| ERR550904 | L5.2 |
| ERR551005 | L5.2 |
| ERR551566 | unknown (new sublineage) |
| ERR551965 | L5.1.1 |
| ERR552187 | L5.1.1 |
| ERR552345 | L5.1.1 |
| ERR702402 | L5.1.1 |
| ERR702411 | L5.1.1 |
| ERR702413 | unknown (new sublineage) |
| ERR702416 | unknown (new sublineage) |
| ERR702417 | L5.1.1 |
| ERR702419 | L5.3 (L5.3.1) |
| ERR702426 | L5.1.1 |
| ERR751295 | L5.1.4 |
| ERR751299 | L5.1.1 |
| ERR751301 | L5.1.1 |
| ERR751302 | L5.1.1 |
| ERR751303 | L5.1.1 |
| ERR751304 | L5.1.3 |
| ERR751310 | L5.1.3 |
| ERR751311 | L5.1.5 |
| ERR751312 | L5.3 (L5.3.1) |

| | |
|---|---|
| ERR751313 | L5.1.1 |
| ERR751315 | L5.1.3 |
| ERR751323 | L5.1.4 |
| ERR751327 | L5.1.1 |
| ERR751328 | L5.1.1 |
| ERR751334 | L5.1.1 |
| ERR751335 | unknown (new sublineage) |
| ERR751339 | L5.1.2 |
| ERR751345 | L5.1.1 |
| 12046-12 | L5.1.5 |
| 12184-03 | unknown (new sublineage) |
| 12910-13 | L5.1.1 |
| 1410-02 | L5.1.3 |
| 1413-02 | L5.1.1 |
| 1426-02 | L5.1.1 |
| 1437-02 | L5.1.1 |
| 1440-02 | L5.1.1 |
| 1465-02 | L5.1.4 |
| 1473-02 | L5.1.4 |
| 14741-14 | L5.1.3 |
| 2568-02 | unknown (new sublineage) |
| 3377-03 | L5.1.1 |
| 3482-03 | L5.1.4 |
| 4005-13 | L5.1.1 |
| 4518-03 | L5.1.1 |
| 4804-03 | L5.1.1 |
| 5378-02 | L5.1.1 |
| 5398-02 | L5.1.1 |
| 5404-02 | L5.1.1 |
| 5405-02 | unknown (new sublineage) |
| 5417-02 | L5.1.2 |
| 5432-02 | L5.1.2 |
| 5434-02 | L5.1.4 |
| 5441-04 | L5.1.1 |
| 5444-02 | L5.2 |
| 5446-02 | unknown (new sublineage) |
| 5447-02 | L5.1.1 |
| 5456-02 | L5.1.1 |
| 5473-02 | L5.1.4 |
| 5475-02 | L5.1.1 |
| 6897-04 | unknown (new sublineage) |
| 8076-11 | L5.2 |
| 8103-11 | L5.2 |
| 8137-11 | L5.2 |
| 8214-03 | L5.1.1 |

| | |
|---|---|
| 8270-03 | L5.1.1 |
| 8303-02 | L5.1.1 |
| 8876-12 | L5.1.1 |
| 8878-12 | L5.1.1 |
| 8946-13 | L5.1.1 |
| 9858-03 | L5.3 (L5.3.1) |
| BSSE-QGF-108892 | L5.1.1 |
| BSSE-QGF-109560 | L5.1.1 |
| BSSE-QGF-109571 | L5.1.1 |
| BSSE-QGF-109627 | L5.1.1 |
| BSSE-QGF-22108 | unknown (new sublineage) |
| BSSE-QGF-22110 | L5.1.3 |
| GR2-CGATGT-L007 | L5.1.1 |
| MTB-DY-135 | L5.1.2 |
| MTB-DY-20 | unknown (new sublineage) |
| MTB-DY-26 | L5.3 (L5.3.1) |
| NG-5288-1047301 | L5.1.1 |
| NG-5288-256902 | L5.1.1 |
| NG-5288-533304 | L5.1.1 |
| NG-5288-553604 | L5.1.1 |
| NG-5967-N1063 | unknown (new sublineage) |
| NG-6193-N1203 | L5.1.1 |
| SRR2100183 | L5.1.2 |
| SRR2100713 | unknown (new sublineage) |
| SRR2101040 | L5.1.2 |
| SRR2101063 | L5.1.2 |
| SRR2101065 | L5.1.4 |
| SRR2101293 | L5.1.1 |
| SRR7496542 | unknown (new sublineage) |
| SRR998585 | L5.1.1 |
| SRR998618 | L5.1.1 |

Table S8. Accession numbers, country of origin and sequencing statistics of Illumina sequenced L5 genomes

| G_NUMBER | AC_NUMBER | Country | Average read depth | Std of read depth | Percentage_not_covered |
|---|---|---|---|---|---|
| G00183 | ERR234142 | Ghana | 80.93 | 39.07 | 2.27% |
| G00188 | ERR234144 | Ghana | 76.54 | 29.9 | 1.65% |
| G00193 | ERR234147 | Ghana | 83.46 | 27.11 | 0.47% |
| G00196 | ERR234149 | Ghana | 87.75 | 28.61 | 0.63% |
| G00410 | ERR841430 | Liberia | 82.95 | 17.69 | 0.81% |
| G00731 | ERR4192047 | Ghana | 62.29 | 14.16 | 0.72% |
| G00829 | ERR841494 | South Africa | 107.03 | 26.57 | 0.70% |
| G00892 | ERR234199 | Ghana | 131.33 | 38.69 | 0.89% |
| G00895 | ERR234202 | Ghana | 32.72 | 11.7 | 1.49% |
| G00898 | ERR234204 | Ghana | 60.94 | 17.86 | 1.39% |
| G01935 | ERR4192364 | Nigeria | 73.63 | 21.26 | 0.69% |
| G01937 | ERR4192366 | Nigeria | 56.86 | 17.92 | 0.91% |
| G01976 | ERR4192367 | Equatorial Guinea | 105.69 | 46.37 | 0.55% |
| G01977 | ERR4192368 | Equatorial Guinea | 89.16 | 21.3 | 0.68% |
| G01978 | ERR4192369 | Equatorial Guinea | 91.39 | 25.77 | 0.57% |
| G01979 | ERR4192370 | Equatorial Guinea | 145.14 | 36.5 | 0.73% |
| G01980 | ERR4192371 | Equatorial Guinea | 61.55 | 30.86 | 0.57% |
| G01981 | ERR4192372 | Equatorial Guinea | 85.46 | 30.78 | 0.69% |
| G01982 | ERR4192373 | Equatorial Guinea | 68.64 | 34.69 | 0.73% |
| G02691 | ERR4192380 | Gambia | 125.46 | 32.27 | 1.23% |
| G03829 | ERR1334053 | Ghana | 96.04 | 17.71 | 0.94% |
| G04163 | ERR1215478 | Ghana | 32.19 | 8.67 | 0.61% |
| G04164 | ERR1215477 | Ghana | 50.23 | 10.28 | 0.58% |

| | | | | | |
|---|---|---|---|---|---|
| G04165 | ERR1215476 | Ghana | 55.72 | 11.26 | 0.60% |
| G04168 | ERR552345 | No Africa | 65.65 | 29.25 | 0.64% |
| G07450 | ERR4192561;ERR4192562;ERR4192563 | Ivory Coast | 86.37 | 26.22 | 0.58% |
| G08166 | ERR4192386;ERR4192407;ERR4192544 | Equatorial Guinea | 61.68 | 20.98 | 0.73% |
| G08388 | ERR1023216 | Gambia | 147.25 | 26.27 | 0.72% |
| G08389 | ERR1023217 | Gambia | 141.73 | 27 | 0.89% |
| G08390 | ERR1023218 | Gambia | 150.36 | 27.06 | 0.69% |
| G08393 | ERR1023221 | Gambia | 154.28 | 31.33 | 1.09% |
| G08395 | ERR1023223 | Gambia | 152.6 | 27.95 | 0.48% |
| G08396 | ERR1023224 | Gambia | 146.32 | 31.4 | 0.64% |
| G08470 | ERR1082117 | Ghana | 177.17 | 34.08 | 0.71% |
| G08472 | ERR1082120 | Ghana | 178.32 | 42.5 | 0.57% |
| G08473 | ERR1082122 | Ghana | 162.75 | 32.02 | 0.54% |
| G08476 | ERR1082125 | Ghana | 176.1 | 46.33 | 0.59% |
| G08477 | ERR1082126 | Ghana | 141.32 | 25.18 | 0.61% |
| G08480 | ERR1082129 | Ghana | 188.23 | 28.52 | 0.56% |
| G08486 | ERR1082135 | Ghana | 151.2 | 25.51 | 0.93% |
| G08488 | ERR1082137 | Ghana | 160.3 | 26.42 | 0.81% |
| G08496 | ERR1203054 | Ghana | 127.36 | 21.51 | 0.77% |
| G08500 | ERR1203057 | Ghana | 119.35 | 22.27 | 0.56% |
| G08501 | ERR1203058 | Ghana | 119.66 | 23.45 | 0.57% |
| G08502 | ERR1203059 | Ghana | 112.83 | 19.47 | 0.59% |
| G08507 | ERR1203065 | Ghana | 112.84 | 18.2 | 0.56% |
| G08508 | ERR1203066 | Ghana | 124.55 | 19.6 | 0.57% |
| G08510 | ERR1203068 | Ghana | 123.36 | 20.14 | 0.52% |
| G08511 | ERR1203069 | Ghana | 112.67 | 20.24 | 0.94% |
| G08516 | ERR1203074 | Ghana | 125.48 | 20.35 | 0.72% |
| G08541 | ERR439931 | Cameroon | 79.93 | 13.46 | 0.05% |
| G08546 | ERR439936 | Guinea | 83.16 | 14.05 | 0.51% |
| G08547 | ERR439937 | Benin | 64.58 | 11.92 | 0.55% |

| | | | | | |
|---|---|---|---|---|---|
| G08549 | ERR439939 | Benin | 57.11 | 11.07 | 0.60% |
| G08550 | ERR439940 | Benin | 65.84 | 12.15 | 0.67% |
| G08551 | ERR439941 | Benin | 67.66 | 11.61 | 0.41% |
| G08554 | ERR439944 | Benin | 74.03 | 12.89 | 0.46% |
| G08557 | ERR439947 | Benin | 64.2 | 11.79 | 0.50% |
| G08559 | ERR439949 | Benin | 65.95 | 12.04 | 0.59% |
| G08561 | ERR439951 | Benin | 78.95 | 14.69 | 1.16% |
| G08562 | ERR439952 | Benin | 75.54 | 13.56 | 0.49% |
| G08563 | ERR439953 | Benin | 70.79 | 12.12 | 0.60% |
| G08565 | ERR439955 | Benin | 66.89 | 12.8 | 1.16% |
| G08569 | ERR439959 | Benin | 60.49 | 13.01 | 1.11% |
| G08570 | ERR439960 | Benin | 58.87 | 10.94 | 0.44% |
| G08572 | ERR439962 | Benin | 62.57 | 11.35 | 0.63% |
| G08574 | ERR439964 | Benin | 64.07 | 12.74 | 0.70% |
| G08577 | ERR439967 | Guinea | 68.33 | 12.44 | 0.55% |
| G08590 | ERR439980 | Ivory Coast | 64.44 | 11.89 | 0.54% |
| G08592 | ERR439982 | Benin | 53.23 | 10.36 | 0.74% |
| G08593 | ERR439983 | Benin | 65.41 | 13.82 | 0.71% |
| G08594 | ERR439984 | Benin | 70.69 | 13.36 | 0.57% |
| G08595 | ERR439985 | Benin | 63.36 | 11.61 | 0.40% |
| G08609 | ERR460916 | Benin | 93.26 | 16.62 | 0.45% |
| G08611 | ERR502471 | Ghana | 79.03 | 13.94 | 0.63% |
| G08612 | ERR502475 | Ghana | 95.84 | 15.31 | 0.17% |
| G08616 | ERR502487 | Ghana | 82.11 | 13.94 | 0.72% |
| G08619 | ERR702402 | Ghana | 76.42 | 14.85 | 0.73% |
| G08622 | ERR502500 | Ghana | 80.33 | 13.65 | 0.69% |
| G08623 | ERR502501 | Ghana | 99.81 | 18.96 | 1.08% |
| G08627 | ERR502505 | Ghana | 89.13 | 17.2 | 0.11% |
| G08631 | ERR502509 | Ghana | 96.96 | 16.51 | 0.52% |
| G08632 | ERR502512 | Ghana | 89.75 | 16.35 | 0.61% |

| | | | | | |
|---|---|---|---|---|---|
| G08633 | ERR502513 | Ghana | 54.44 | 14.2 | 0.92% |
| G08635 | ERR502515 | Ghana | 80.8 | 15.83 | 0.66% |
| G08652 | ERR502536 | Ghana | 77.92 | 14.2 | 0.91% |
| G08656 | ERR1215473 | Ghana | 47.9 | 11.37 | 0.55% |
| G08660 | ERR702407 | Ghana | 74.28 | 14.65 | 0.54% |
| G08664 | ERR702411 | Ghana | 76.88 | 15.2 | 0.69% |
| G08666 | ERR702413 | Ghana | 75.3 | 12.3 | 0.52% |
| G08669 | ERR702416 | Ghana | 75.09 | 16.34 | 0.56% |
| G08670 | ERR702417 | Ghana | 80.41 | 15.1 | 0.54% |
| G08672 | ERR702419 | Ghana | 77.66 | 12.49 | 0.51% |
| G08679 | ERR702426 | Ghana | 74.15 | 12.52 | 0.54% |
| G08695 | ERR751295 | Ghana | 86.88 | 17.5 | 0.92% |
| G08699 | ERR751299 | Ghana | 84.94 | 16.75 | 0.56% |
| G08701 | ERR751301 | Ghana | 81.22 | 14.65 | 0.54% |
| G08702 | ERR751302 | Ghana | 87.89 | 16.87 | 0.70% |
| G08703 | ERR751303 | Ghana | 83.14 | 15.78 | 0.54% |
| G08704 | ERR751304 | Ghana | 92.78 | 18.52 | 0.74% |
| G08708 | ERR751308 | Ghana | 77.59 | 13.77 | 0.39% |
| G08710 | ERR751310 | Ghana | 55.2 | 16.93 | 0.65% |
| G08711 | ERR751311 | Ghana | 80.87 | 14.44 | 0.50% |
| G08712 | ERR751312 | Ghana | 83.18 | 16.82 | 0.38% |
| G08713 | ERR751313 | Ghana | 77.85 | 15.8 | 0.60% |
| G08715 | ERR751315 | Ghana | 82.84 | 13.98 | 0.55% |
| G08721 | ERR751323 | Ghana | 89.55 | 14.95 | 0.68% |
| G08723 | ERR751327 | Ghana | 67.08 | 11.91 | 0.56% |
| G08724 | ERR751328 | Ghana | 65.72 | 12.13 | 0.56% |
| G08730 | ERR751334 | Ghana | 86.4 | 16.54 | 0.60% |
| G08731 | ERR751335 | Ghana | 75.31 | 13.26 | 0.64% |
| G08735 | ERR751339 | Ghana | 49.43 | 9.68 | 0.58% |
| G08741 | ERR751345 | Ghana | 69.72 | 12.98 | 0.54% |

| | | | | | |
|---|---|---|---|---|---|
| G08743 | ERR751348 | Ghana | 92.05 | 16.94 | 0.91% |
| G08848 | ERR1215463 | Ghana | 44.99 | 9.31 | 0.52% |
| G08913 | ERR019875 | No Africa | 29.23 | 11.6 | 1.64% |
| G08917 | ERR234679 | No Africa | 126.4 | 20.86 | 0.85% |
| G08918 | ERR234680 | No Africa | 116.97 | 19.1 | 0.51% |
| G08920 | ERR551566 | Sierra Leone | 64.74 | 17.79 | 0.77% |
| G08934 | SRR998585 | Mali | 151.95 | 32.17 | 0.12% |
| G11003 | SRR998618 | Mali | 151.66 | 36.31 | 0.32% |
| G11556 | ERR3170399 | Ghana | 52.65 | 14.83 | 0.86% |
| G11565 | ERR3170440 | Ghana | 86.9 | 21.54 | 0.77% |
| G11566 | ERR3170441 | Ghana | 44.99 | 13.94 | 1.19% |
| G11568 | ERR4162017 | Gabon | 59.41 | 16.96 | 0.74% |
| G11575 | ERR551089 | Sierra Leone | 66.53 | 19.82 | 1.12% |
| G11578 | ERR3170443 | Ghana | 67.25 | 17.96 | 0.71% |
| G11579 | ERR551857 | Sierra Leone | 61.23 | 17.31 | 0.76% |
| G11591 | ERR3170456 | Ghana | 105.34 | 25.94 | 0.66% |
| G11595 | ERR3170460 | Ghana | 45.89 | 12.79 | 1.01% |
| G11597 | ERR3170463 | Ghana | 42.7 | 12.71 | 0.70% |
| G11598 | ERR3170464 | Ghana | 122.54 | 34.01 | 0.85% |
| G11600 | ERR3170466 | Ghana | 113.82 | 36.95 | 0.77% |
| G11604 | ERR3170471 | Ghana | 47.59 | 13.62 | 1.02% |
| G11605 | ERR552588 | Ghana | 73.32 | 20.22 | 1.06% |
| G11607 | ERR3170473 | Ghana | 49.4 | 13.09 | 1.02% |
| G11608 | ERR3170474 | Ghana | 91.38 | 23.93 | 0.69% |
| G11609 | ERR3170475 | Ghana | 51.37 | 16.02 | 0.76% |
| G11610 | ERR3170476 | Ghana | 65.15 | 18.85 | 0.71% |
| G11611 | ERR3170477 | Ghana | 43.03 | 11.35 | 0.72% |
| G11614 | ERR3170480 | Ghana | 67.64 | 23.17 | 0.83% |
| G11619 | ERR3170483 | Ghana | 71.16 | 19.26 | 0.58% |
| G11620 | ERR3170484 | Ghana | 105.94 | 26.91 | 0.75% |

| | | | | | |
|---|---|---|---|---|---|
| G11626 | ERR3170490 | Ghana | 104.41 | 24.65 | 0.67% |
| G11627 | ERR3170491 | Ghana | 49.54 | 13.68 | 0.73% |
| G11631 | ERR4162025 | Gabon | 56.05 | 15.47 | 0.90% |
| G11632 | ERR4162026 | Gabon | 102.51 | 29.46 | 0.85% |
| G11633 | ERR4162027 | Gabon | 117.26 | 31.58 | 0.68% |
| G11637 | ERR3148046 | Ghana | 81.18 | 21 | 0.50% |
| G11639 | ERR4162032 | Democratic Republic of the Congo | 81.46 | 20.79 | 0.63% |
| G11643 | ERR3170404 | Ghana | 104.28 | 28.22 | 0.96% |
| G11646 | ERR3170407 | Ghana | 49.53 | 15.96 | 0.82% |
| G11649 | ERR552187 | Ghana | 68.92 | 19.83 | 0.96% |
| G11654 | ERR3170413 | Ghana | 75.04 | 20.49 | 0.87% |
| G11655 | ERR4162000 | Gabon | 75.66 | 20.02 | 0.88% |
| G11656 | ERR4162001 | Gabon | 83.32 | 23.6 | 0.56% |
| G11657 | ERR4162002 | Gabon | 133.3 | 35.31 | 0.72% |
| G11658 | ERR551336 | Sierra Leone | 64.78 | 17.85 | 0.78% |
| G11659 | ERS4575215 | Gabon | 95.25 | 23.79 | 0.83% |
| G11660 | ERR4162005 | Gabon | 79.69 | 21.03 | 0.88% |
| G11661 | ERR3170414 | Ghana | 56.8 | 14.98 | 0.97% |
| G11663 | ERR4162006 | Gabon | 97.1 | 27.08 | 0.69% |
| G11666 | ERR3170417 | Ghana | 66.83 | 16.98 | 0.78% |
| G11667 | ERR3170418 | Ghana | 64.88 | 16.11 | 0.80% |
| G11668 | ERR3170419 | Ghana | 89.74 | 23.14 | 0.71% |
| G11671 | ERR3170422 | Ghana | 117.23 | 28.65 | 0.65% |
| G11672 | ERR3170423 | Ghana | 104.37 | 28.35 | 0.72% |
| G11674 | ERR3170424 | Ghana | 117.45 | 32.15 | 0.72% |
| G11679 | ERR3170428 | Ghana | 47.74 | 13.31 | 1.15% |
| G11683 | ERR552506 | Ghana | 57.64 | 18.39 | 1.16% |
| G11685 | ERR4162007 | Gabon | 119.87 | 32.72 | 0.80% |
| G11693 | ERR3170434 | Ghana | 175.53 | 43.2 | 0.58% |
| G11696 | ERR552261 | Sierra Leone | 63.95 | 18.67 | 0.85% |

| | | | | | |
|---|---|---|---|---|---|
| G11698 | ERR3148058 | Gabon | 167.21 | 46.01 | 0.67% |
| G11700 | ERR551005 | Republic of the Congo | 69.42 | 17.32 | 0.74% |
| G11701 | ERR551124 | Republic of the Congo | 69.8 | 17.67 | 0.85% |
| G11702 | ERR552673 | Republic of the Congo | 64.88 | 18.67 | 1.15% |
| G11703 | ERR550904 | Republic of the Congo | 122.71 | 31.58 | 0.67% |
| G11704 | ERR552285 | Republic of the Congo | 64.04 | 16.89 | 0.68% |
| G11706 | ERR3170488 | Ghana | 90.82 | 29.02 | 0.61% |
| G14330 | SRR2100183 | No Africa | 65.57 | 12.1 | 0.54% |
| G14855 | SRR2100713 | No Africa | 51.05 | 12.71 | 0.33% |
| G15178 | SRR2101040 | No Africa | 68.07 | 13.78 | 0.47% |
| G15201 | SRR2101063 | No Africa | 55.9 | 13.93 | 0.75% |
| G15203 | SRR2101065 | No Africa | 51.53 | 11.67 | 0.68% |
| G15430 | SRR2101293 | No Africa | 55.26 | 12.58 | 0.62% |
| G22215 | ERR2704692 | Ghana | 55.29 | 14.56 | 0.63% |
| G25357 | ERR550904 | Republic of the Congo | 122.71 | 31.58 | 0.67% |
| G26731 | SRR11444083 | Ghana | 55.19 | 11.16 | 0.63% |
| G26738 | SRR11444026 | Ghana | 47.69 | 9.74 | 0.67% |
| G26749 | SRR11444232 | Ghana | 62.94 | 10.52 | 0.42% |
| G26805 | SRR11444206 | Ghana | 69.52 | 12.03 | 0.55% |
| G32940 | ERR2383619 | Cameroon | 115.01 | 28.88 | 0.81% |
| G32941 | ERR2383620 | Cameroon | 149.11 | 35.82 | 0.79% |
| G32942 | ERR2383621 | Ivory Coast | 89.61 | 20.8 | 0.57% |
| G32943 | ERR2383622 | No Africa | 82.08 | 21.12 | 0.51% |
| G32944 | ERR2383623 | No Africa | 69.76 | 18.79 | 0.88% |
| G32945 | ERR2383624 | No Africa | 83.77 | 22.73 | 0.86% |
| G32946 | ERR2383625 | No Africa | 92.66 | 24.9 | 0.72% |
| G32947 | ERR2383626 | No Africa | 82.86 | 20.7 | 0.54% |
| G32950 | ERR2704808 | Ivory Coast | 327.61 | 42.04 | 0.60% |
| G32951 | ERR2704809 | Cameroon | 316.26 | 40.72 | 0.58% |
| G32952 | ERR2704810 | Cameroon | 329.55 | 40.39 | 0.41% |

| | | | | | |
|---|---|---|---|---|---|
| G32954 | ERR2704812 | Cameroon | 378.7 | 59.3 | 1.35% |
| G33338 | SRR7496542 | No Africa | 146.46 | 25.1 | 0.46% |
